# Distinct epigenetic and metabolic states reflect the regional identity of adult neural stem cells and prime their fate

**DOI:** 10.64898/2026.01.26.701741

**Authors:** Valentina Scandella, Nikolaos Lykoskoufis, Hadrien Soldati, Livia Di Martino, Tony Teav, Hector Gallart-Ayala, Julijana Ivanisevic, Simon M.G. Braun, Marlen Knobloch

## Abstract

Neural stem/progenitor cells (NSPCs) in the adult mouse brain reside in distinct niches, including the dentate gyrus (DG) and subventricular zone (SVZ). The contribution of cell intrinsic versus extrinsic factors to distinct fates of NSPCs and their neuronal progeny remains largely unknown. We here show that DG- and SVZ-derived NSPCs retain niche-specific chromatin accessibility states that predict neuronal subtype specification. Furthermore, metabolic profiling and gene expression analyses comparing DG- and SVZ-derived NSPCs revealed differences in carnitine synthesis pathways and lipid metabolism. Supplementation with carnitine or S-adenosylmethionine (SAM) induced chromatin remodeling via histone modifications, indicating that a metabolic-epigenetic cross-talk determines regional neuronal identity. Our findings show that NSPCs are intrinsically programmed by chromatin and metabolic states, and that metabolic interventions alter epigenetic landscapes and fate potential.

## Introduction

Neural stem/progenitor cells (NSPCs) generate new neurons throughout life in a process called adult neurogenesis, which occurs in specific neurogenic niches. In rodents, two main neurogenic niches have been identified: the subventricular zone (SVZ) lining the lateral ventricle and the subgranular zone (SGZ) of the hippocampal dentate gyrus (DG) (*1–3*). Neurogenic niches contain various cell types that sustain and influence the whole neurogenic process, from exiting quiescence to proliferation, fate decisions and neuronal differentiation (*4*, *5*). NSPCs of the SGZ and SVZ are both radial-glia-like cells that can self-renew, proliferate and give rise to differentiated progeny. They express common molecular markers and rely on conserved neurogenic signaling pathways, highlighting their close functional and developmental relationship (*6–12*).

However, there is a fundamental difference in the type and function of neurons they generate. In the SVZ, NSPCs give rise to neuroblasts, which migrate along the rostral migratory stream (RMS) to reach the olfactory bulb (OB) and differentiate into a variety of olfactory interneurons. These mainly inhibitory neurons integrate into the olfactory circuitry, playing roles in olfactory discrimination and plasticity, and contribute to tissue homeostasis (*13–19*). In the DG, neuroblasts differentiate into excitatory neurons that integrate into the local hippocampal circuit and contribute to memory formation, pattern separation, and hippocampus-dependent plasticity (*20–26*). How much these differences are due to intrinsic differences in the two NSPC populations or instructed by their niche is poorly understood. Several studies have used transplantation approaches to probe for cell-intrinsic versus niche-derived signals that regulate fate and behavior of NSPCs and their progeny (*27–29*). Adult NSPCs from DG transplanted to the RMS, or SVZ NSPCs transplanted into the hippocampus, have shown that adult NSPCs have a flexibility to adapt to the region they are transplanted to, supporting the concept that the local microenvironment is a strong contributor to NSPC behavior (*27*). Conversely, other transplantation studies showed that the regional identity of NSPCs influences their grafting and integration capacity, highlighting that there is also a strong contribution of cell-intrinsic mechanisms (*28*, *29*).

Furthermore, to understand what regulates adult neurogenesis, many studies profiled and perturbed molecular pathways in adult NSPCs and their progeny, revealing important mechanisms, including epigenetic regulatory processes (*30–35*), gene- and protein-expression differences along fate change (*36–44*), as well as specific metabolic pathways which influenced NSPC behavior (*45–52*). However, the majority of those studies has been done in a niche-specific manner and comparative studies on molecular characteristics done in parallel between SVZ and DG NSPCs are limited (*53*).

To determine the extent to which cell-intrinsic differences contribute to the distinct fate of adult NSPCs, we isolated and cultured SVZ and DG NSPCs simultaneously from individual mice, and performed a thorough analysis of their epigenetic, gene expression, and metabolic states. Our data show that NSPCs display distinct chromatin and metabolic signatures according to their regional identity, which prime them to generate specific types of neurons in a cell-intrinsic way. Furthermore, region-specific differences in gene expression were maintained throughout neural differentiation *in vitro*, and modulating cellular metabolism of NSPCs perturbed these differences, suggesting a metabolic-epigenetic crosstalk which influences fate potential.

## Results

### Individually cultured DG and SVZ NSPCs proliferate in a similar manner

In order to investigate whether DG- and SVZ-derived adult NSPCs retain their regional identity outside of their respective niches when cultured *in vitro*, we sub-dissected the hippocampus and the SVZ from individual 8-week-old C57BL/6J wildtype male and female mice to isolate and propagate DG- and SVZ-derived NSPCs simultaneously (Fig. 1A). To enrich for a population of stem cells, NSPC cultures were kept as neurospheres for at least four consecutive passages (more than 25 days *in vitro*) prior to the analysis (Fig. 1A).

**Figure 1:**
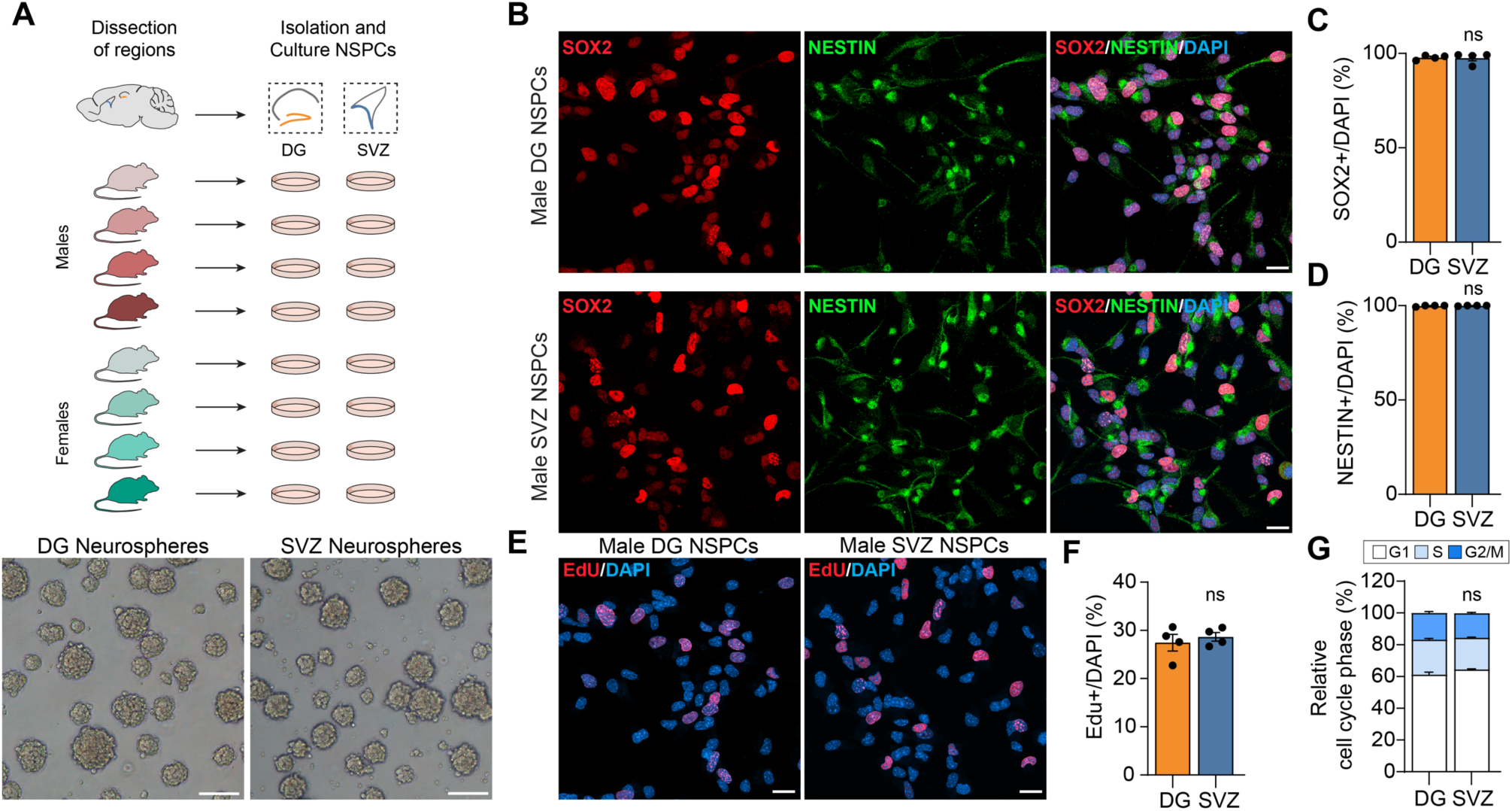
Male DG and SVZ NSPCs harbor a similar proliferation profile. **A)** Experimental outline of cultured DG and SVZ NSPCs isolated from 4 male and 4 female adult mice. Bottom panel shows brightfield image of neurospheres cultured for 1 month. Scale bar: 100 µm. **B)** Immunofluorescence staining for SOX2 and NESTIN in DG and SVZ male NSPCs. Scale bar: 20 µm. **C)** Quantification of SOX2^+^ cells in DG and SVZ male NSPCs. Data show individual replicates and mean ± SEM. N= 4 biological replicates, unpaired Student t-test, ns = not significant. **D)** Quantification of NESTIN^+^ cells in DG and SVZ male NSPCs. Data show individual replicates and mean ± SEM. N= 4 biological replicates, unpaired t-test: ns. **E)** Immunofluorescence staining for EdU in DG and SVZ male NSPCs. Scale bar: 20 µm. **F)** Quantification of EdU^+^ cells in DG and SVZ male NSPCs. Data show individual replicates and mean ± SEM. N=4 biological replicates, unpaired t-test. ns **G)** Quantification of DG and SVZ NSPC cell cycle stages by flow cytometry. Data show mean ± SEM. N= 3 biological replicates, unpaired t-test, ns.

To confirm the presence of NSPCs, for each individual culture we quantified the number of cells expressing two stem cells markers, the nuclear marker SRY-Box transcription factor 2 (SOX2) and the filament protein Nestin. Nearly 100% of male and female NSPCs from SVZ and DG expressed both markers (Fig. 1B-D, Fig. S1A, B). We assessed proliferation using a one-hour 5-ethynyl-2’-deoxyuridine (EdU) labeling, allowing the detection of cells that underwent S-phase within this time. About 30% of NSPCs incorporated EdU, with no difference between SVZ and DG in both male and female NSPCs (Fig. 1E, F, Fig. S1C). Cell-cycle profiling using flow cytometry in pooled DG and SVZ male NSPCs confirmed a similar cell cycle profile (Fig. 1G).

Taken together, these data show that NSPC cultures from individual male and female mice from DG and SVZ have a similar proliferation rate and marker expression, regardless of the sex and region of origin.

### Despite prolonged culture in vitro, DG and SVZ NSPCs keep a distinct chromatin accessibility profile, which predicts the type of neurons they will generate

We next assessed whether *in vitro* culture and the absence of niche signals influenced the regional identity of DG and SVZ NSPCs. We used a genome-wide Assay for Transposase-Accessible Chromatin (ATAC-seq) to profile the epigenetic state of chromatin(*54*) on all 16 individual NSPC cultures (Fig. 2A). Principle component analysis (PCA) showed that the original region was the main driver of clustering samples together, while sex played a less important role (Suppl. Fig. 2A). We thus used sex chromosomes as covariates for further analysis, to be able to pool the data from individual male and female samples from the same neurogenic region for downstream analysis (n=8 per region). Indeed, the PCA of the ATAC-seq data showed that DG and SVZ NSPCs retain a specific chromatin accessibility profile even when cultured *in vitro* (Fig. 2B).

**Figure 2:**
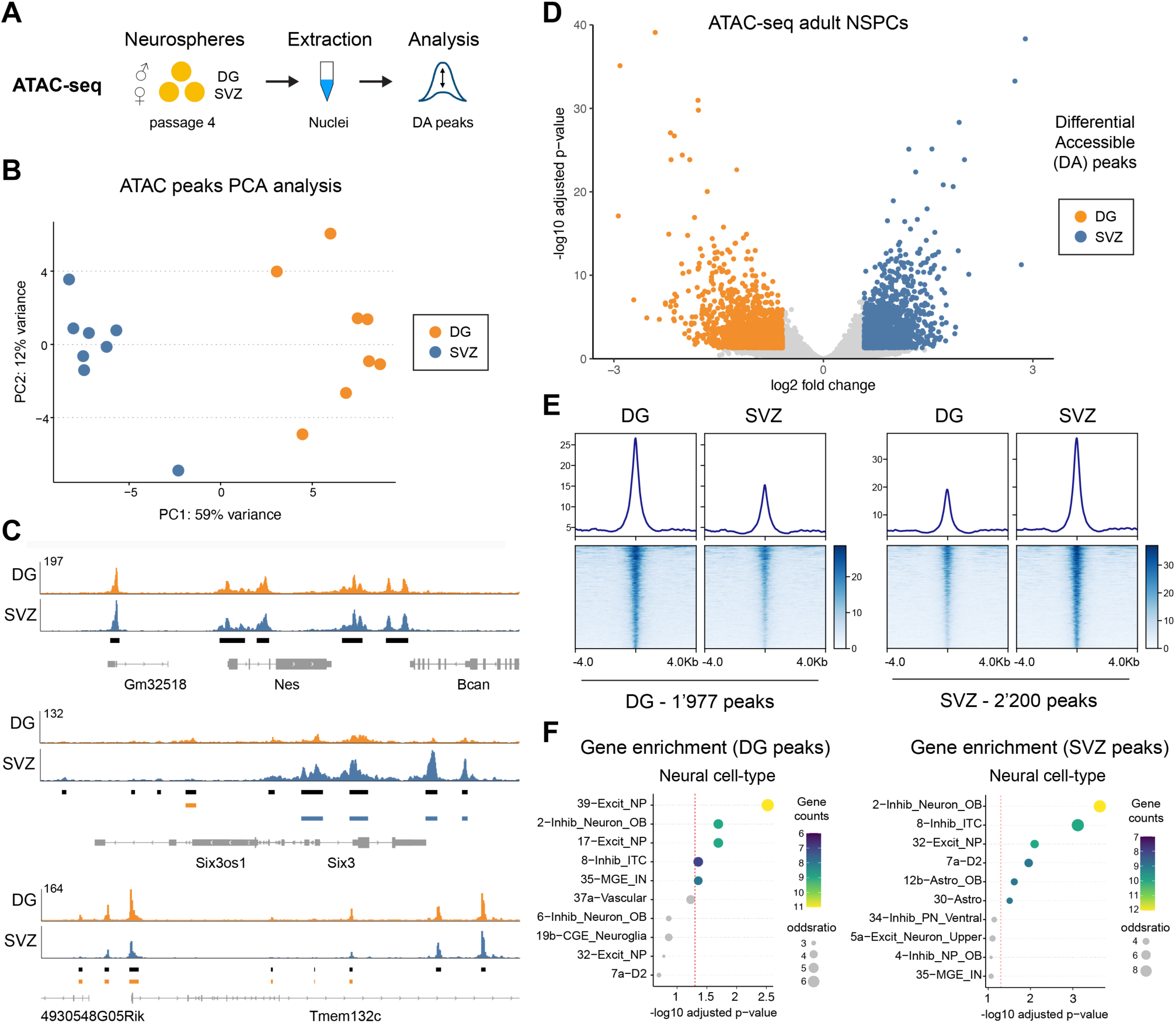
ATAC-seq analysis of DG and SVZ NSPCs predicts the type of neurons they generate. **A)** Experimental scheme of ATAC-seq analysis in cultured DG and SVZ NSPCs. **B)** PCA of ATAC-seq data from DG (orange) and SVZ (blue) NSPCs. N= 8 biological replicates. **C)** Genome browser tracks of ATAC-seq peaks at the *Nestin, Six3 and Tmem132c* loci for DG NSPCs (orange) and SVZ NSPCs (blue). N = 8 biological replicates, average bigwig file displayed. **D)** Volcano plot of differential accessible peaks between DG and SVZ NSPCs. The orange dots represent regions more accessible in DG NSPCs and blue dots represent more accessible regions in SVZ NSPCs. FDR < 0.05, FC > |1.5|. N = 8 biological replicates. **E)** ATAC-seq signal in DG and SVZ NSPCs at DG and SVZ specific peaks, centered at the peak center and ranked according to read intensity. **F)** Neural cell-type enrichment analysis of the genes associated with DG NSPC-specific ATAC peaks (left) and SVZ NSPC-specific ATAC peaks (right).

Among the 132’412 peaks identified by ATAC-seq, 4’177 regions showed significant differences in accessibility between DG and SVZ (Suppl. Table 1). This suggests that while the two cell types have overall similar chromatin accessibility profiles, the SVZ- and DG-specific ATAC peaks may contribute to genetic programs that control the future cell fate of these progenitors (Fig. 2C-E, Suppl. Fig. 2B). Notably, the differentially accessible genomic regions were enriched for enhancers rather than promoters, which are regulatory elements enriched with cell-type specific TF binding motifs that control gene expression networks during development. (Suppl. Fig. 2C-D).

To better understand the differences in chromatin accessibility between DG and SVZ NSPCs, we further examined the differential accessible DA peaks and identified the genes associated with these changed peaks. As one of the main differences between DG and SVZ NSPCs *in vivo* is the type of progeny they give rise to, namely excitatory neurons for DG NSPCs and mainly inhibitory interneurons for SVZ NSPCs, we questioned whether the DA peaks would reflect this difference. We therefore looked for a signature of prospective neurons the two types of NSPCs generate. We used a recently established list of 53 brain cell type clusters (as measured by scRNA-seq (*55*)) and compared it to our list of genes associated with DA peaks. Surprisingly, in the DG NSPCs, genes associated with DG-specific ATAC peaks were enriched for genes associated with the cluster of “excitatory neuronal progenitors (NP)” (Fig. 2F). On the contrary, genes associated with DA peaks in SVZ NSPCs were enriched for genes characteristic of inhibitory neurons of the OB (Fig. 2F). Strikingly, the cell types identified by ATAC-Seq data analysis exactly match the neuronal subtypes that NSPCs in the DG and SVZ will produce once they are differentiating, suggesting that the fate of these cells is indeed encoded at an epigenetic level and maintained *in vitro*.

### The differential neuronal fate priming is not detected at the gene expression level in cultured NSPCs

Chromatin accessibility can directly reflect gene expression, with open regions correlating with increased expression and more close regions with gene silencing. However, many changes in chromatin accessibility can also be non-transcriptional, and only influence gene expression at a later stage, for instance during differentiation(*56*, *57*). To determine if this fate priming was already evident at the gene expression level, we performed bulk RNA barcoding and sequencing (BRB-seq)(*58*) of all 16 NSPC cultures (Fig. 3A). Similarly to the ATAC-seq data, PCA showed that the original region was the main driver of sample clustering, while sex played a less important role (Suppl. Fig. 3A). We thus again used the sex chromosomes as covariates for downstream analysis. The PCA showed a clear separation of the 16 samples according to their region of origin (Fig. 3B). Analysis of the differentially expressed genes (DEGs) between DG and SVZ NSPCs showed a similar number of upregulated (1095) and downregulated (859) genes (Fig. 3C), of which around one third (28%) overlapped with the DA peaks (Fig. 3D and Suppl. Fig. 3B, Suppl. Table 2). As all the cultures were established and expanded individually, we used an unpaired analysis, but even when taking the original pairing, (meaning the mouse from which the SVZ and DG NSPCs were extracted) into account, we obtained highly similar lists of DEGs (>98% overlap) (Suppl. Fig. 3C and D).

**Figure 3:**
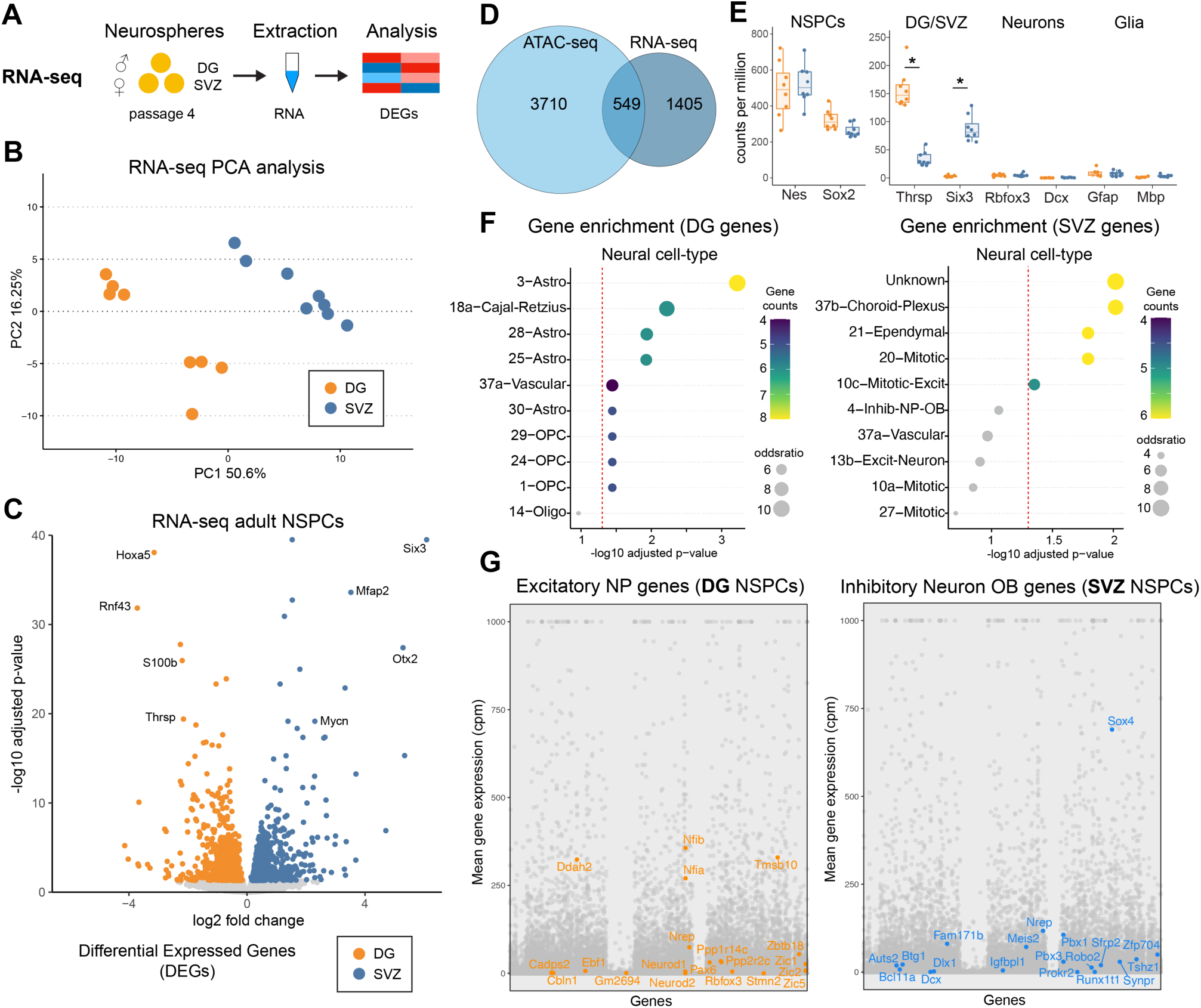
RNA-seq analysis of DG and SVZ NSPCs reveals no changes in expression of neuronal fate priming genes. **A)** Experimental scheme of RNA-seq analysis in cultured DG and SVZ NSPCs. **B)** PCA of RNA-seq data from DG (orange) and SVZ (blue) NSPCs. N= 8 biological replicates. **C)** Volcano plot of differential gene expression between DG and SVZ NSPCs. The orange dots represent genes more expressed in DG NSPCs and blue dots represent genes more expressed in SVZ NSPCs. FDR < 0.05. N = 8 biological replicates. **D)** Venn diagram displaying the overlap between differentially expressed genes (RNA-seq) and genes associated with differential accessible peaks (ATAC-seq). **E)** Normalized RNA-seq counts for DG and SVZ NSPCs at marker genes for NSPCs (*Nes, Sox2)*, DG/SVZ NSPCs (*Thrsp, Six3*), neurons (*Rbfox3, Dcx*) and glia (*Gfap, Mbp*). N = 8 biological replicates. Unpaired t-test, *adj. p-value <0.05. **F)** Neural cell-type enrichment analysis of the differentially expressed genes in DG NSPCs (left) and SVZ NSPCs (right). **G)** Dot plot representing mean gene expression level for all genes in DG NSPCs (left) and SVZ NSPCs (right). Labelled in orange are the genes that characterize the “excitatory neuronal progenitor” cell-type. Labelled in blue are the genes that characterize the “inhibitory neuron of the olfactory bulb” cell-type.

We next performed the same cell type analysis as for the ATAC-seq data using the DEGs of the RNA-seq analysis. As seen with immunohistochemical approaches, classical NSPC markers such as *Nestin* and *Sox2* were similarly expressed, whereas differentiation markers for neurons (*Rbfox3*) and glial cells (*Gfap* and *Mbp*) were very lowly expressed and not different between the two NSPC types (Fig. 3E). However, genes that have been previously described to differ between NSPC types such as thyroid hormone-inducible hepatic protein (*Thrsp*) and Homeobox protein SIX3 (*Six3*) were also differentially expressed between DG and SVZ NSPCs in culture (Fig. 3E). In contrast to the ATAC-seq, the DEGs did not predict a distinct neuronal type. The cell type “NSPC” was not present among the 53 cell types in the published scRNA-seq atlas clusters used for this analysis, and therefore the cell type “astrocyte” was most enriched for DG NSPCs (Fig. 3F), whereas for SVZ NSPCs the cell types “Choroid plexus” and “Ependymal cells” were the top matches (Fig. 3F). As expected, the genes that were used to determine the cell type “excitatory NP and “inhibitory OB neurons” were only very lowly expressed and at similar levels between DG and SVZ NSPCs (Fig. 3G and Suppl. Fig. 3E). Thus, these data suggest that while NSPCs in culture retain a distinct gene expression pattern, matching their regional origin, they are still in an undifferentiated state and have not already changed their gene expression towards a specific neuronal fate. Thus, the specific “fate-priming” seen in the ATAC-seq data seems to be transcriptionally silent and might only become expressed under differentiation conditions. This suggests that the differentiation potential of NSPCs is encoded in the chromatin accessibility landscape, most notably at enhancers, before the expression of specific markers of neuronal subtypes.

### NSPCs from the two niches are metabolically distinct despite prolonged culture in vitro

Recent studies have revealed a dynamic interplay between cellular metabolism and epigenetic modifications, as certain metabolites are directly required for histone modifications(*59–64*). We thus assessed if differences in chromatin accessibility may also be reflected by distinct metabolic states. We extracted the intracellular metabolites of each of the individual 16 NSPC cultures and performed untargeted profiling using liquid chromatography (LC) coupled with high resolution mass spectrometry (HRMS) assay (Fig. 4A). Within the acquired multiparametric profiles we could annotate a portion of polar metabolome (n=134 metabolites) with high confidence. These annotated 134 metabolites were used for the following analyses. The principal component analysis (PCA) of NSPC metabolic profiles revealed that the neurogenic region the NSPCs were extracted from, namely DG and SVZ, determined the sample clustering, whereas sex played only a minor role (Fig. 4B). Similar to the RNA-seq and ATAC-seq data, we therefore decided to combine male and female samples for subsequent statistical analyses. To better evaluate the specific metabolic signature of DG and SVZ NSPCs, we looked into differences at the metabolite level. 14 metabolites were significantly more abundant in DG NSPCs and 21 metabolites were present in higher levels in SVZ NSPCs (Fig. 4C).

**Figure 4:**
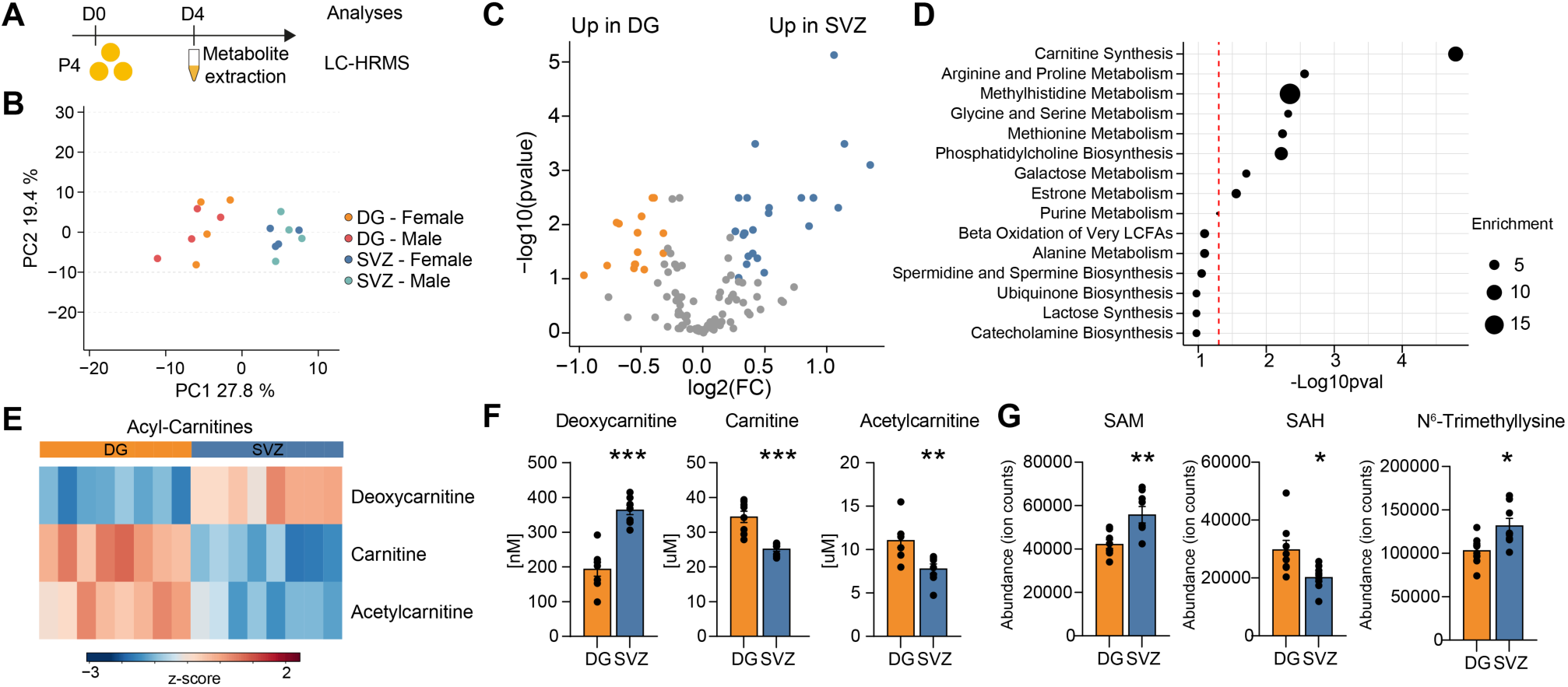
Metabolic profiling of DG and SVZ NSPCs reveals differences in the carnitine synthesis pathway. **A)** Experimental outline of metabolomics profiling of cultured DG and SVZ NSPCs. **B)** PCA of metabolic signatures from male and female DG NSPCs (orange) and SVZ NSPCs (blue), acquired using untargeted metabolomics approach. N= 8 biological replicates. **C)** Volcano plot of identified metabolites in DG and SVZ NSPCs. The orange dots represent metabolites with higher abundance in DG NSPCs and blue dots represent metabolites with higher abundance in SVZ NSPCs. Cutoffs: adjusted p-value =0.1, fold change of 20%. N= 8 biological replicates. **D)** Pathway enrichment analysis of the significant metabolites (adjusted p-value 0.1). Red dotted line represents p-value =0.05, circle size represents enrichment of the pathway. **E)** Heatmap of three significantly changed metabolites involved in the carnitine synthesis pathway in DG and SVZ NSPCs, p-value=0.05. **F)** Quantification of carnitine synthesis-related metabolites. Data show individual replicates and mean ± SEM. N=8 biological replicates. Unpaired t-test, * p<0.05, ** p<0.01, *** p<0.001 **G)** Relative abundance (in ion counts) of selected metabolites S-adenosylmethionine (SAM), S-adenosylhomocysteine (SAH) and ^6^N-Trimethyllysine from untargeted metabolomics. Data show individual replicates and mean ± SEM. N=8 biological replicates. Unpaired t-test, corrected for multiple comparisons, * p<0.05, ** p<0.01.

### DG and SVZ NSPCs show differences in the carnitine synthesis pathway

A metabolite set enrichment analysis (MSEA) using the quantitative enrichment analysis (QEA) showed that the most significantly different pathway was carnitine synthesis (Fig. 4D). To corroborate these findings, we performed a comprehensive targeted quantification of carnitine and acylcarnitines. This analysis yielded the concentrations for 28 acylcarnitines measurable in NSPCs derived from the SVZ and DG (Table 1). The precursor of carnitine, deoxycarnitine (also known as butyrobetaine), was higher in the SVZ NSPCs compared to DG NSPCs (Fig. 4E and F, Table 1). Interestingly, carnitine and its acetylated form, acetylcarnitine, were found in higher amounts in DG NSPCs (Fig. 4E and F). In addition, among all quantified carnitines species, a few others showed a significant difference between DG and SVZ, yet with a less clear pattern between DG and SVZ NSPCs (Suppl. Fig. 4A and B, Table 1). Together, these data suggest that there are clear differences in carnitine levels between DG and SVZ NSPCs, despite culture in medium containing the same nutrients.

**Table 1:** Absolute quantification of carnitine species in DG and SVZ NSPCs.

| Metabolites | Unit | DG |  | SVZ |  | Statistics |  |
| --- | --- | --- | --- | --- | --- | --- | --- |
|  |  | Average | SEM | Average | SEM | pval | padj |
| Stearoylcarnitine | nM | 17.692 | 3.921 | 48.123 | 19.468 | 0.1661 | 0.301 |
| Oleoylcarnitine | nM | 46.781 | 8.202 | 102.487 | 23.916 | 0.0563 | 0.142 |
| Octadecadienoylcarnitine | nM | 3.046 | 0.613 | 6.552 | 1.718 | 0.0878 | 0.196 |
| Palmitoylcarnitine | nM | 31.995 | 7.538 | 49.799 | 12.407 | 0.2445 | 0.373 |
| Hexadecenoylcarnitine | nM | 16.616 | 2.915 | 31.754 | 7.648 | 0.0974 | 0.202 |
| Myristoylcarnitine | nM | 21.796 | 4.528 | 26.485 | 5.578 | 0.5250 | 0.609 |
| Tetradecenoylcarnitine | nM | 6.441 | 1.007 | 13.400 | 2.942 | 0.0533 | 0.142 |
| Tetradecanedi enoylcarnitine | nM | 0.898 | 0.176 | 2.018 | 0.485 | 0.0588 | 0.142 |
| Lauroylcarnitine | nM | 4.778 | 0.969 | 5.960 | 1.113 | 0.4368 | 0.528 |
| Dodecenoylcarnitine | nM | 0.815 | 0.195 | 2.174 | 0.563 | 0.0493 | 0.142 |
| Decanoylcarnitine | nM | 1.808 | 0.385 | 2.362 | 0.415 | 0.3447 | 0.476 |
| Octanoylcarnitine | nM | 2.682 | 0.613 | 2.880 | 0.518 | 0.8084 | 0.808 |
| Hydroxyoctadecanoylcarnitine | nM | 3.574 | 0.949 | 6.672 | 2.522 | 0.2802 | 0.406 |
| Hexanoylcarnitine | nM | 17.233 | 2.981 | 16.069 | 1.837 | 0.7454 | 0.772 |
| Hydroxyhexadecanoylcarnitine | nM | 4.294 | 0.560 | 7.246 | 2.054 | 0.2028 | 0.346 |
| Hexenoylcarnitine | nM | 4.398 | 2.317 | 3.584 | 0.697 | 0.7448 | 0.772 |
| Hydroxytetradecanoylcarnitine | nM | 3.204 | 0.471 | 7.250 | 1.722 | 0.0530 | 0.142 |
| Isovalerylcarnitine | nM | 58.887 | 7.210 | 108.824 | 10.235 | 0.0016 | 0.016 |
| Hydroxydodecanoylcarnitine | nM | 2.053 | 0.357 | 3.655 | 1.222 | 0.2429 | 0.373 |
| Tiglylcarnitine | nM | 6.241 | 0.458 | 10.495 | 0.982 | 0.0029 | 0.021 |
| Butyrylcarnitine | nM | 1331.466 | 115.492 | 973.472 | 93.462 | 0.0310 | 0.128 |
| Propionylcarnitine | nM | 478.521 | 38.007 | 395.496 | 33.101 | 0.1222 | 0.236 |
| Deoxycarnitine | nM | 193.582 | 20.058 | 364.256 | 13.283 | 0.0000 | 0.000 |
| Hydroxyhexanoylcarnitine | nM | 35.841 | 4.193 | 33.554 | 4.562 | 0.7176 | 0.772 |
| Acetylcarnitine | uM | 11.047 | 0.789 | 7.795 | 0.541 | 0.0051 | 0.029 |
| Hydroxyvalerylcarnitine | nM | 1.887 | 0.469 | 2.562 | 0.544 | 0.3635 | 0.479 |
| Hydroxybutyrylcarnitine | nM | 558.349 | 66.283 | 635.441 | 65.713 | 0.4227 | 0.528 |
| Carnitine | nM | 34.429 | 1.612 | 25.159 | 0.625 | 0.0004 | 0.006 |

Carnitine is essential for lipid metabolism: it enables the transport of activated fatty acids (acyl-CoAs) into mitochondria for fatty acid β-oxidation (FAO)(*65*), a pathway utilized by proliferating SVZ and DG NSPCs, and which is also implicated in NSPC maintenance (*45*, *51*, *52*). Carnitine synthesis requires ^6^N-trimethylated lysine (TML), which is a post-translational amino acid modification involving S-adenosylmethionine (SAM) and lysine methyltransferases (*66*). TML is released after the degradation of methylated proteins like histones and is considered a rate-limiting factor for carnitine synthesis (*67*).

We thus re-examined carnitine pathway-related metabolites from the untargeted analysis dataset (Fig. 4G) and found that TML and S-adenosylmethionine (SAM) were present in significantly higher levels in SVZ NSPCs compared to DG NSPCs, whereas the level of S-adenosylhomocysteine (SAH) was increased in DG NSPCs (Fig. 4G).

Taken together, these results identify that many intermediates in the carnitine pathway are significantly altered between NSPCs from DG and SVZ.

### Lipid metabolism related genes are also altered in ATAC-seq and RNA-seq datasets

We next assessed whether the metabolic differences were also reflected at the chromatin accessibility and gene expression level. As there was no obvious difference in the genes involved in carnitine biosynthesis (Suppl. Table 2.), we performed a more global pathway enrichment analysis with a curated list of metabolism-related genes for each of the datasets (DA peaks and DEGs). Genes that showed increased chromatin accessibility or gene expression in DG samples pointed indeed towards lipid metabolism (metabolism of lipids, fatty acid metabolism, triglyceride metabolism) (Fig. 5A, Suppl. Fig. 5A). On the other hand, genes with increased accessibility and expression in SVZ were associated with RNA metabolism and amino acids (Fig. 5A and Suppl. Fig. 5B). Together, these data show that the metabolic differences are at least in part regulated at the chromatin accessibility and gene expression levels.

**Figure 5:**
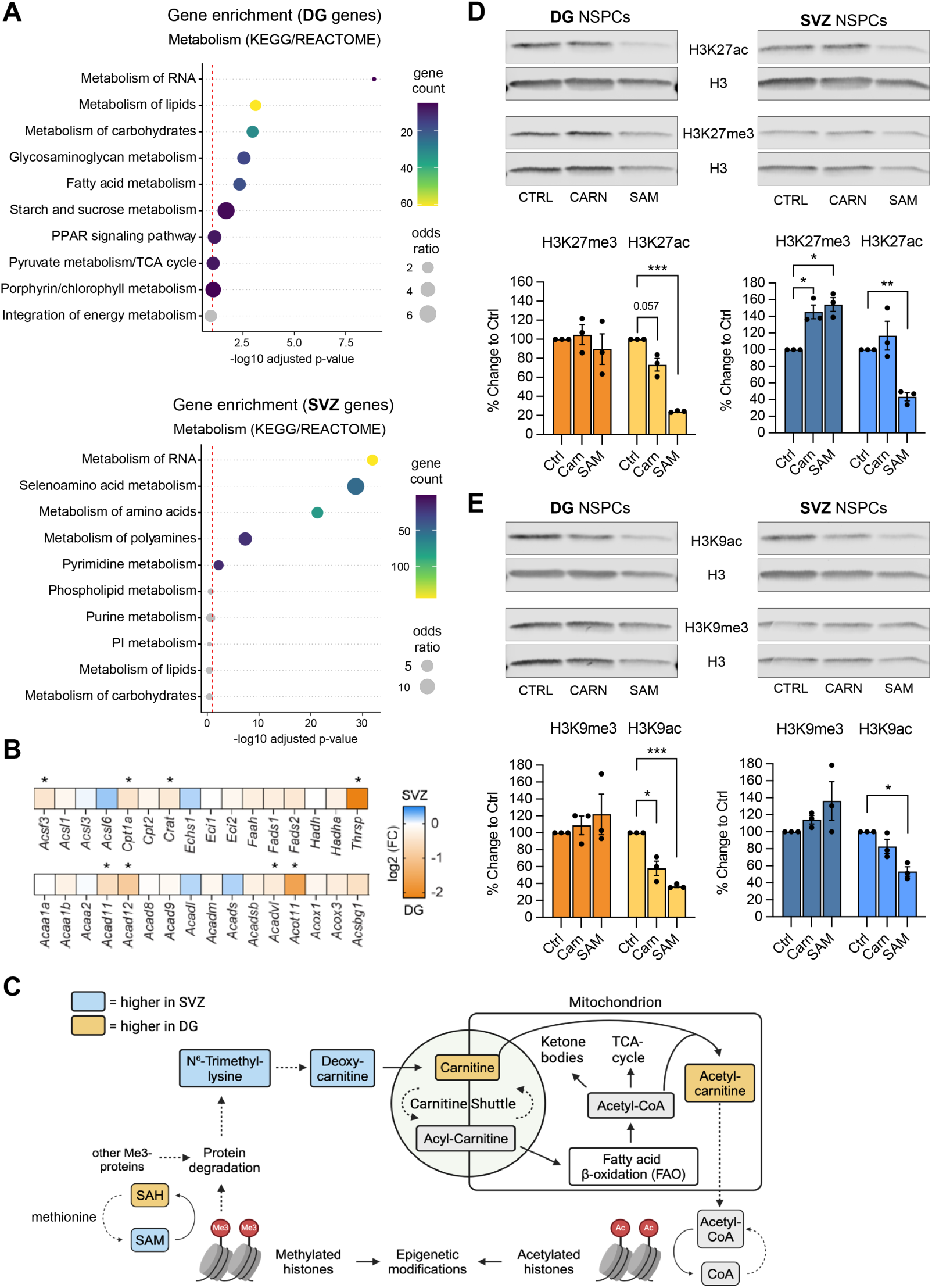
Supplementation with carnitine or SAM alters histone acetylation and methylation levels in DG and SVZ NSPCs. **A)** KEGG/REACTOM pathway enrichment analysis of the upregulated DEGs in DG NSPCs (top) and in SVZ NSPCs (bottom), red dotted line represents p=0.05. **B)** Heatmap of FAO-related genes. Color-coded according to log2 fold change, orange labels genes more expressed in DG NSPCs and blue more expressed in SVZ NSPCs. *adjusted p-value <0.05 and log2 (FC) ± 0.3. **C)** Schematic representation of links between S-adenosylmethionine (SAM), acetyl-CoA and carnitine metabolism with colours representing levels of individual metabolites between DG and SVZ NSPCs. **D and E)** Western blot analysis of histone acetylation (H3K27ac/H3K9ac) and histone methylation (H3K27me3/H3K9me3) levels normalized to total histone H3 levels in control (CTRL) DG and SVZ NSPCs, as well as NSPCs treated with carnitine (CARN) or SAM for 5 days. N = 3 samples per condition. One-sample t-tests for treatment effect. * p<0.05, ** p<0.01, *** p<0.001. Of note, cell numbers were lower after SAM treatment, which is also reflected in reduced total H3 levels.

The observed changes point towards altered lipid metabolic pathways. As carnitine is essential for FAO, we next looked in more details at the gene expression of genes involved in FAO. Indeed, many FAO genes were upregulated in DG NSPCs compared to SVZ NSPCs (Fig. 5B), among which the rate limiting enzyme *Cpt1a,* as well as acyl-CoA dehydrogenases (*Acad11-12*), very long-chain acyl-CoA dehydrogenase (*Acadvl*) and acyl-CoA thioesterase 11 (*Acot11*). In addition, *Crat*, which catalyzes the transfer of acetyl groups from coenzyme A to carnitine, is also higher in DG samples (Fig. 5B). Of note, *Thrsp*, which has been shown to be a specific DG NSPC marker (*46*), is strongly upregulated in DG NSPCs compared to SVZ NSPCs (Fig. 5B). Taken together, these data suggest that DG NSPCs have a higher transport of FAs into mitochondria and do more FAO.

### Carnitine metabolism is linked to epigenetic modifications in NSPCs and supplementation with carnitine or SAM affects their epigenetic profiles

Epigenetic regulation and carnitine biosynthesis are directly linked through histone methylation (*66*). Indeed, TML is an important post-translationally modified amino acid found on histones and it is necessary for carnitine synthesis. Furthermore, carnitine is needed for FAO, which breaks down fatty acids into acetyl-CoA. This acetyl-CoA can then be used in the tricarboxylic acid (TCA) cycle for energy production or exported from the mitochondria via carnitine for other use, including for histone acetylation (Fig. 5C). Given the differences between DG and SVZ NSPCs in these pathways at the metabolic, gene expression and chromatin accessibility level (Fig. 5C), we next explored whether there were differences at the histone modification level between DG and SVZ NSPCs. Additionally, we investigated whether supplementing the cell culture medium with carnitine or SAM, two metabolites that exhibited significant differences in DG and SVZ and serve as sources of acetyl-CoA and methyl-donors respectively, could result in changes in these epigenetic marks. For this, we pooled the individual cultures of male NSPCs from each region, treated them for 5 days with carnitine (1mM) or SAM (0.1mM), extracted cell nuclei and performed western blot analyses of different histone methylation and acetylation marks (Fig. 5D and E). We focused our analysis on marks associated with active chromatin (H3K4me3, H3K27ac and H3K9ac) and with repressive chromatin (H3K27me3. H3K9me3). We could not detect significant baseline differences in these epigenetic marks between DG and SVZ NSPCs, which might be due to the limited sensitivity of western blot analysis (Suppl. Fig. 5C and D). However, both treatments with carnitine and SAM lead to significant changes in several histone modifications (Fig. 5D and E). Indeed, we observed an increase in global H3K9me3 and H3K27me3 levels upon treatment with SAM in both DG and SVZ NSPCs. These SAM-induced changes coincided with global losses in H3K9ac and H3K27ac. Furthermore, carnitine treatment displayed similar changes in histone acetylation and methylation in NSPCs, albeit at more modest levels (Fig. 5D and E). These data reveal that key drivers of chromatin state, H3K9me3/ac and H3K27me3/ac, are directly regulated by cellular metabolism, as exogenous treatment with carnitine or SAM greatly impact global levels of these epigenetic marks in NSPCs.

### The regional identity is maintained during differentiation

All the differences described so far point towards a distinct regional identity, which is maintained even when NSPCs are kept for a prolonged time in culture and which is in part influenced by their metabolic status. Furthermore, our ATAC-seq data strongly suggest that this regional identity will predict future neuronal fate, although the NSPCs are in a stem cell state and do not yet express differentiation markers. However, whether these changes have indeed a functional consequence on the actual differentiation into neurons is not clear. We therefore performed an *in vitro* differentiation experiment by withdrawing growth factors from the medium, thereby triggering spontaneous differentiation into neurons and astrocytes. To do so, we used pooled NSPCs from individual cultures of male DG and SVZ NSPCs, plated them on laminin and collected samples for RNA-seq at day 0, day 2, day 4 and day 6 (Fig. 6A). PCA analysis showed that the changes which occur upon differentiation were driving the clustering of the samples, with the samples clustering according to the trajectory of differentiation, with d6 samples being furthest apart from the d0 samples (Fig. 6B). As expected, NSPC and proliferation markers went down over the course of differentiation in both populations, whereas neuronal and glial markers went up (Fig. 6C and D, Suppl. Fig. 6A and Suppl. Table 3). Interestingly, the neuronal marker *Dcx*, which is expressed by migrating neuroblasts and immature neurons, came up earlier in the differentiating SVZ NSPCs compared to DG NSPCs, whereas the more mature marker *Rbfox3* (encoding for NeuN) came up earlier in DG NSPCs (Fig. 6C). These differences might correlate with the known differences in migratory behavior, as progenitor cells from the SVZ migrate a long distance as neuroblasts (DCX-positive) to the olfactory bulb, whereas progenitor cells from the DG mature locally.

**Figure 6:**
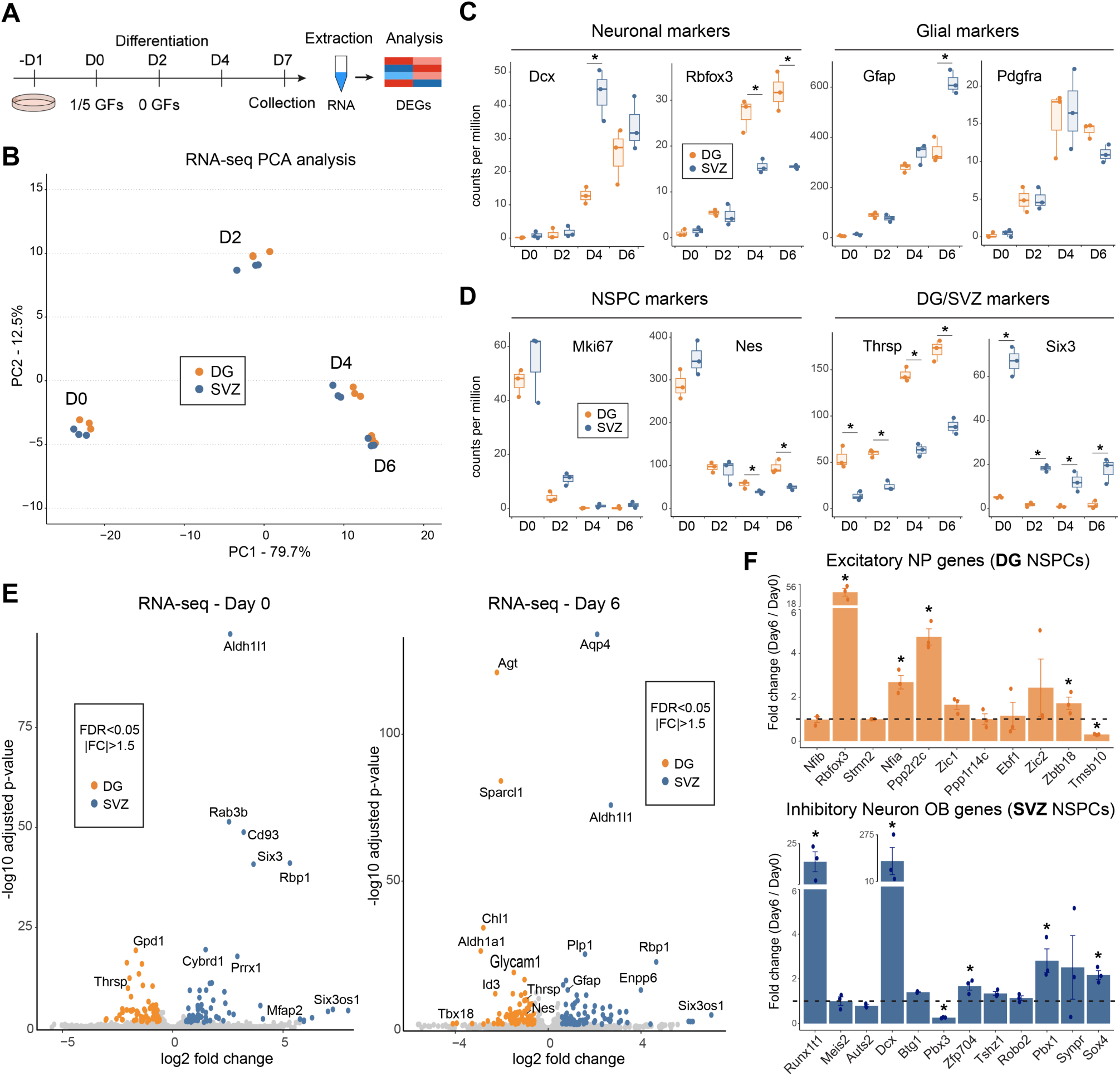
The regional identity of DG and SVZ NSPCs is maintained during differentiation. **A)** Timeline of differentiation experiment in DG and SVZ NSPCs. **B)** PCA of RNA-seq data from DG NSPCs (orange) and SVZ NSPCs (blue) at Day 0, 2, 4 and 6 of differentiation. N= 3 biological replicates per timepoint and cell type. **C)** Normalized RNA-seq counts for DG and SVZ NSPCs across indicated differentiation timepoints for marker genes for neurons (*Dcx, Rbfox3*) and glia (*Gfap, Pdgfra*). **D)** Normalized RNA-seq counts for DG and SVZ NSPCs across indicated differentiation timepoints for marker genes for NSPCs (*Mki67, Nes)* and DG/SVZ markers (*Thrsp, Six3*). N = 3 biological replicates. Unpaired t-test, * adj. p<0.05. **E)** Volcano plots of differential gene expression between DG and SVZ NSPCs at Day 0 and Day 6 of differentiation. FDR < 0.05 and FC > |1.5|. N = 3 biological replicates per timepoint and cell type. **F)** Graphs represent fold-change between Day 0 and Day 6 for genes that characterize the “excitatory neuronal progenitor” cell-type in DG NSPCs, and genes that characterize “inhibitory neuron of the olfactory bulb” cell-type in SVZ NSPCs.

In line with our previously identified DEGs, genes that contributed to the regional identity were also differentially expressed at d0 (Fig. 6D and E, Suppl. Fig. 6A). Interestingly, dozens of DEGs were maintained throughout the differentiation process and were still apparent at d6 (Fig. 6E and Suppl. Fig. 6A and B). Finally, we found that certain marker genes identified by ATAC-seq as predicting the neuronal fate of DG and SVZ NSPCs were indeed expressed in day 6 neurons (Fig. 6F). For example, while not expressed in DG and SVZ NSPCs at day 0, *Rbfox3* and *Ppp2r2c* were upregulated in DG neurons at d4 and d6, whereas *Runx1t1* and *Dcx* were higher in SVZ neurons at d4 and d6 (Suppl. Fig. 6C and D). Taken together, these RNA-seq data highlight that, as in the adult mouse brain, DG and SVZ NSPCs differentiate into distinct population of neurons *in vitro* despite lacking niche signals.

### Carnitine and SAM treatment of NSPCs influences gene expression profiles during differentiation

As carnitine and SAM treatment of NSPCs had led to epigenetic changes (Fig. 5D and E), we wanted to assess whether this treatment prior to differentiation could also influence gene expression during differentiation and whether this would influence their regional identity profile. As before, pooled NSPCs from male DG and SVZ were treated for 5 days with Carnitine or SAM under proliferation condition, then plated for differentiation, without Carnitine or SAM, and samples were collected at d0, d2, d4 and d6 and analysed by RNA-seq (Fig. 7A). PCA analysis showed that the differentiation was still the main driver of clustering, despite the different pre-treatments (Suppl. Fig. 7A). When looking at the regional identity genes, it became apparent that the pre-treatment with either carnitine or SAM led to a disturbance in the expression of these genes, as shown with fold change heatmaps for DEGs across conditions and timepoints (Fig. 7B). Whereas the separation of the DG and SVZ NSPCs was clear throughout differentiation in control conditions, numerous genes displayed changes in relative expression in DG and SVZ cells upon carnitine or SAM treatment (Fig. 7B). For example, *Tbx2*, *Rbm46, Six3*, and *Col9a1,* markers of DG and SVZ cells in control cells were no longer as strongly differentially expressed following carnitine or SAM treatments (Fig. 7C and 7D). Similarly, expression of the differentiation markers *Dcx* and *Rbfox3* were also modestly altered, although not reaching statistical significance (Suppl. Fig. 7B and C). This analysis suggests that modulation of the carnitine pathway through carnitine or SAM supplementation alters the epigenetic mechanisms that control DG and SVZ identity gene expression during differentiation. Taken together, these results suggest that adult neural stem cells can maintain their regional identity and differential potential outside of their respective niches, via the metabolic regulation of their epigenetic states.

**Figure 7:**
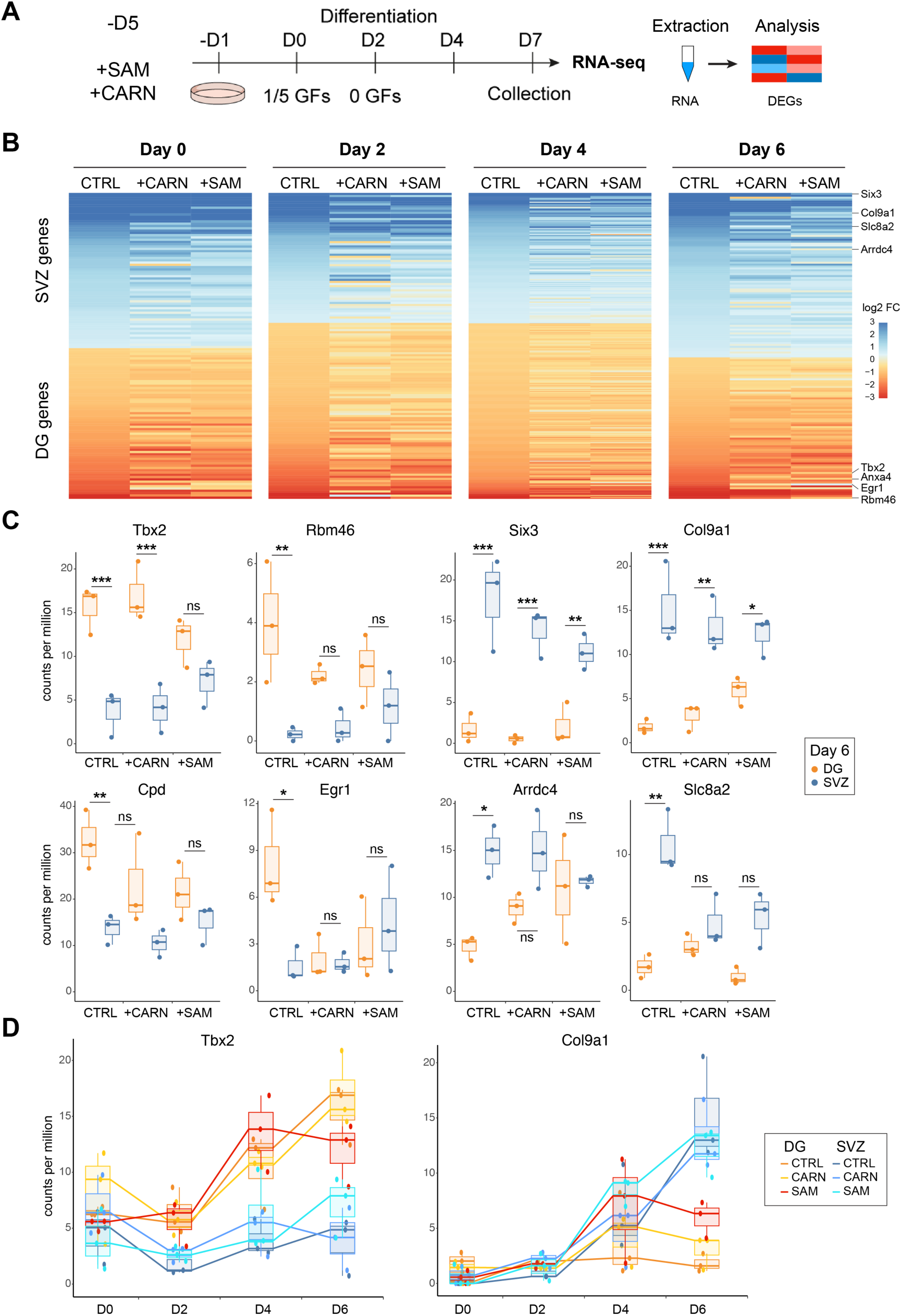
Carnitine and SAM treatment of NSPCs influences expression of neuronal-identity genes during differentiation. **A)** Experimental scheme of RNA-seq analysis in control (CTRL) DG and SVZ NSPCs, as well as NSPCs treated with carnitine (CARN) or with S-adenosylmethionine (SAM) for 5 days before induction of differentiation for a further 6 days. **B)** Heatmaps represent log2 fold change of DG and SVZ specific genes in CTRL, CARN and SAM treated conditions across all timepoints. **C)** Normalized RNA-seq counts for DG and SVZ NSPCs at Day 6 of differentiation in CTRL, CARN and SAM treated conditions. Shown are some of the most differentially expressed genes between DG and SVZ cells under CTRL condition (see heatmap in B), which change with CARN or SAM treatment (*Tbx2, Rbm46, Six3, Col9a1, Cpd, Egr1, Arrdc4, Slc8a2*). N = 3 biological replicates per timepoint, cell type and treatment. Unpaired t-test, * adj. p<0.05, ** adj. p<0.01, *** adj. p<0.001. ns, non-significant. **D)** Normalized RNA-seq counts for DG and SVZ cells across all differentiation timepoint in CTRL, CARN and SAM treated conditions, for two marker genes of DG and SVZ cells (*Tbx2, Col9a1*). N = 3 biological replicates per timepoint, cell type and treatment.

## Discussion

The ability of stem cells to receive and integrate a large variety of niche signals to control their proliferation and fate decisions has been described for many different stem cell niches, including the two adult neurogenic niches, the DG and SVZ (*2*, *3*). However, as stem cells also use cell-intrinsic mechanisms to control their fate, and given the complexity of the niches, it is challenging to disentangle the detailed mechanisms. In this study, we used a simplified approach to address the cell-autonomous contribution of NSPC fate regulation by isolating NSPCs simultaneously from the DG and SVZ from individual mice and expanding them *in vitro* under the same growth conditions to remove potential influences by niche factors. We performed systematic analyses of gene expression (RNA-seq) and chromatin accessibility (ATAC-seq), as well as metabolic profiling. We found a remarkable difference based on the region of origin in both male and female NSPCs, which was kept despite prolonged culture *in vitro*. Chromatin dynamics is an important regulator of NSPC behavior: several studies have found differentially accessible genomic loci between quiescent/active NSPCs as well as young/old NSPCs (*68–71*). These chromatin states are established by epigenetic regulators and are crucial for NSPC fate decisions, especially during developmental neurogenesis, where the NSPCs (called radial glia, RG) undergo drastic fate changes from early neurogenic to later gliogenic (*72*). For example, the Polycomb Repressive Complex 2 (PRC2) that places repressive H3K27me3 modifications, is a key regulator of RG activity, as PRC2 deletion results in accelerated AP differentiation into neurons (*73*).

However, direct comparisons of ATAC-seq profiles between adult DG and SVZ NSPCs have not been described so far. Remarkably, chromatin accessibility data obtained here clearly show that genes important for the future progeny of NSPCs, namely olfactory bulb interneurons for SVZ NSPCs and excitatory neurons for DG NSPCs, were already primed through differential chromatin accessibility at regulatory sequences. These data point towards a strongly preserved epigenetic fate priming based on the regional identity of adult NSPCs, which is kept even under *in vitro* culture conditions, meaning in a niche-independent manner. This is in line with previous studies that showed that adult SVZ NSPCs retain the expression of specific transcription factors or generate specific subtypes of neurons with regard to their more dorsal or ventral location in the brain (*29*, *74*).

Interestingly, the observed differences in chromatin accessibility do not seem to influence the NSPCs under proliferation conditions, as only a few genes that predicted fate on the ATAC-seq level were robustly expressed at the RNA level. This is in line with the concept that chromatin accessibility can be transcriptionally silent and only influence gene expression under specific conditions, as is the case upon differentiation (*56*). Furthermore, expressed NSPC genes showed high accessibility within their promoters for both cell types, however the differences in chromatin accessiblity bewteen DG and SVZ NSPCs were enriched in enhancer regions, regulatory sequences that control developmental gene expression (*75*).

Over the last decade, it has become clear that cellular metabolism not only provides energy and macromolecules for the cell to function, but can also directly influence gene expression, for instance through providing metabolites for histone acetylation or methylation, which in turn influence chromatin accessibility (*62*, *72*, *76*). To address whether differences in metabolism could explain the observed differences in chromatin accessibility between DG and SVZ NSPCs, we performed untargeted and targeted metabolomics. Our approach to simultaneously isolate and expand NSPCs from the DG and SVZ as individual cultures from individual mice allowed us to address in an unbiased manner to what extent the original identity of the NSPCs was influencing their metabolic profile. Surprisingly, despite *in vitro* culture, NSPCs kept a distinct metabolic profile, although they displayed identical stemness characteristics and proliferation rates. The carnitine synthesis pathway was the main contributor to the divergent clustering. Carnitine is required for the transport of long chain fatty acids into mitochondria for further breakdown via FAO(*65*). Carnitine and acetylcarnitine were higher in DG NSPCs, suggesting an increase in the transport of FAs into mitochondria. In line with these changes, expression of FAO related genes suggested an increase in FAO in DG NSPCs compared to SVZ NSPCs. Both SVZ and DG NSPCs have been previously described to use FAO but were not directly compared (*45*, *51*, *52*). Our data point towards differences in FAO between the two NSPC types, at least *in vitro*. However, whether these differences are kept *in vivo* and whether they depend on nutrient availability remains to be determined. Our metabolomic analyses showed that the differences in carnitine metabolism were not only one-sided: While carnitine and acetylcarnitine were significantly increased in DG NSPCs, the immediate precursor of carnitine, deoxycarnitine, was higher in SVZ NSPCs, as was TML, its original precursor. Differences in the enzyme that converts deoxycarnitine to carnitine, γ-Butyrobetaine hydroxylase (also known as BBOX1) could explain such a difference, however expression levels of *Bbox1* in proliferating NSPCs where too low to be detected in our RNA-seq analysis. An increased metabolic demand of DG NSPCs for high carnitine levels to sustain FAO might lead to an increased efficiency in producing carnitine or an increase in carnitine uptake, however, the exact reasons for the carnitine differences remain to be determined.

Our findings that chromatin accessibility, gene expression and metabolic profiles remain distinct between DG and SVZ NSPCs raises the question of how these differences are maintained in the absence of niche signals. The concept of metabolic-epigenetic crosstalk in this context is intriguing. This concept refers to the bidirectional interaction whereby cellular metabolism provides metabolites that can be used for epigenetic modifications, which in turn alter the expression of metabolic genes (*72*). This can create a feedback loop which could stabilize a certain cellular state. In this regard, the carnitine synthesis pathway is an interesting candidate in two ways: first, carnitine sustains the production of FAO-derived acetyl-CoA, which can be used for histone acetylation, which in turn can directly influence gene expression of lipid metabolism related genes (*64*, *77*). Second, it also requires TML for its synthesis, which is generated when methylated lysines are freed through protein degradation, and methylated histones are an important source of TML (*66*). Thus, carnitine is linked to both histone methylation and histone acetylation. We could however not detect baseline differences in specific histone methylation and acetylation marks between DG and SVZ NSPCs, which might be due to the limited sensitivity of the western blot analysis. Other methods such as spike-in ChIP-seq or CUT&RUN would be needed to determine which specific modifications and which specific genomic loci are differentially regulated by this metabolic process. Nevertheless, treatment of the NSPCs with either SAM or carnitine had a significant effect on the tri-methylation and acetylation of H3K9 and H3K27. As expected, SAM treatment increased histone methylation levels across the genome and resulted in lower histone acetylation, which can be explained by a competition for the same lysine substrate (*78*). Carnitine supplementation did also alter both epigenetic marks. While we would have expected to see an increase in histone acetylation, as other studies in cancer cells have shown (*79*, *80*), we rather observed a decrease in histone acetylation and a modest increase in histone methylation, suggesting that the mechanism is not straight forward through simply increasing FAO and acetyl-CoA. Furthermore, to understand how global methyl-donor and acetyl-CoA levels regulate specific histone modifications at specific genomic loci, future studies focusing on the many different histone modifiers that add this layer of specificity are required. Thus, our data provide a first insight into this metabolic-epigenetic crosstalk in DG and SVZ NSPCs and paves the way for future studies to address these remaining open questions.

A key difference between the two NSPC populations is the progeny they generate, namely excitatory neurons for DG NSPCs and olfactory interneurons for SVZ NSPCs (*2*, *3*). Therefore, a straightforward approach to test the robustness of the epigenetic fate priming would be to differentiate NSPCs and see whether their fates change through metabolic manipulation. However, spontaneous *in vitro* differentiation of NSPCs does not lead to fully mature neurons over the time course of 7 days and neurons cannot be kept much longer without additional supporting factors under these conditions. Furthermore, *in vivo* it takes 4-6 weeks for the progeny to fully mature into the corresponding neuronal type (*21*). Thus, we were technically limited to a less mature neuronal stage. Nevertheless, the robustness of the distinct regional identities was also evident during spontaneous differentiation: although both populations differentiated into neurons and astrocytes with similar morphology and classic marker gene expression (GFAP and DCX), dozens of genes that distinguished DG and SVZ NSPCs under proliferation conditions were also kept distinct throughout the differentiation time course. In line with a pre-determined fate, certain markers that distinguish olfactory bulb neurons and glutamatergic neurons were also clearly differentially expressed between the two NSPC-derived neurons, suggesting that a prolonged and optimized neuronal differentiation would indeed lead to the predicted progeny type. Previous studies have shown that epigenetic manipulation, affecting for instance histone acetylation or methylation in adult NSPCs can indeed influence their fate (*30*, *81*). Here, we find that pretreatment of DG and SVZ NSPCs with two metabolites, carnitine and SAM, perturbs the distinct differentiation profile. These data suggest that the distinct fates are at least in part maintained through a metabolic-epigenetic crosstalk, which is independent of niche signals. Thus our results underline the importance of cellular metabolism for NSPC fate regulation, even under a controlled *in vitro* setting. How different the two niches are *in vivo* in terms of metabolism and how this influences the fate priming remains to be determined.

In summary, our findings highlight the interplay between chromatin accessibility, metabolism, and epigenetic regulation in maintaining NSPC identity and guiding differentiation. The identification of regional signatures suggests that core features of adult NSPCs are intrinsically programmed, and that metabolic interventions can modulate their epigenetic landscape and fate potential.

## Author contributions

V.S. designed, performed, analyzed and interpreted experiments. N.L. performed ATAC and BRB-seq analyses. H.S. performed western blot analyses, T.T. and H.G-A. performed and analyzed metabolomics data with V.S. J.I. designed and interpreted metabolomics data. L.DM performed experiments. S.M.G.B. designed, analyzed and interpreted ATAC and BRB seq data. S.M.G.B. and M.K. developed the concept, designed, performed, analyzed and interpreted experiments. V.S., S.M.G.B. and M.K wrote the manuscript with input from all authors. All authors have seen and approved the final version of the manuscript.

## Acknowledgements

We would like to thank Vanille Maillard for technical help. We also thank the Cellular Imaging facility and the Metabolomics & Lipidomics Platform at University of Lausanne, and the IGE3 Sequencing platform at University of Geneva for technical help. We are further grateful to Sebastian Jessberger for helpful comments on the manuscript. The laboratory of S.M.G.B. is supported by the University of Geneva and the Swiss National Science Foundation (SNSF #PCEFP3_194305), the laboratory of M.K. is supported by the University of Lausanne and the Swiss National Science Foundation.

## Methods

### Animals

C57BL/6 male and female mice used for cell isolation were bought from Janvier (France). All studies were approved by the local authorities (Cantonal Veterinary Office, Canton de Vaud, Switzerland).

### Cell culture

Adult mouse neural stem/progenitor cells (NSPCs) from the SVZ and hippocampi of 7 week-old C57/Bl6 male and female mice were isolated as previously described (*47*). Briefly, after a brief anesthesia, mice were decapitated, and hippocampus and SVZ were sub-dissected. A single-cell suspension per animal and region using a papain-based MACS Neural Tissue Dissociation Kit (#130-092-628, Milteny) was prepared and myelin was removed using the MACS myelin removal beads (#130-096-733, Milteny). The resulting cell preparations were cultured as neurospheres in DMEM/F12 GlutaMAX (#31331-028, Invitrogen) supplemented with B27 (#17504044, Invitrogen), human EGF (20 ng/ml, #AF-100-15, Peprotech), human FGF-basic (20 ng/ml, #100-18B, Peprotech) and PSF (#15240062, Invitrogen), called hereafter B27 medium.

For experiments with adherent NSPCs on coated plates, heparin (5 mg/ml) was added to the proliferative B27 medium. Medium was changed every 2-3 days. NSPCs were used for experiments from passage 4 till passage 10.

#### Proliferation condition

Proliferative NSPCs were grown in B27 medium as described here above. For metabolomics, ATAC-seq and BRB-seq experiments, 200’000 cells were plated on a 6-well format and cultured as neurospheres. Medium was changed every second day, and cells were collected after 4-5 days in culture. Cells were spun down at 100g, cell pellets were washed with PBS and spun again at 300g. Supernatant was sucked off, and cell pellets were snap-frozen in dry-ice and stored at -80° C prior to analysis.

For imaging, 42’000 cells/cm^2^ proliferative NSPCs were plated on coated glass coverslips (10337423, Fisher) with Poly-L-ornithine (50 μg/ml, #P36655, Sigma) and laminin (5 μg/ml, #L2020, Sigma) in a B27 medium containing 5 mg/ml of heparin. 5-ethynyl-2′-deoxyuridine (EdU) pulse was performed by incubating NSPCs with 10 μM EdU for 1 hour at 37°C before fixation.

For cell cycle analysis, proliferative NSPCs cultured in proliferation medium were seeded on coated plastic plate (31’000 cells/cm^2^) for two days.

#### Treatment conditions and cell collection

For the experiments including Carnitine and SAM treatements, cells from the 4 individual SVZ and DG cultures from male mice were pooled per region, taking the same cell numbers per mouse. Cells were expanded as pooled cells and stocks were frozen for subsequent experiments.

For Western blot analyses of the histone marks, 750’000 cells were plated in 6-well plates with 5ml proliferation medium and where either not treated (Ctrl), treated with 1mM L-Carnitine (Sigma-Aldrich #C0158, diluted in ddH2O) or 0.1mM SAM (Sigma-Aldrich #A4377). Medium was changed on the 3rd day, with fresh L-Carnitine and SAM for the treated conditions. Spheres were collected at day 5 as described above. For Western Blots, cells were collected either without HDAC inhibitor for the methylation marks, or with 1x HDAC inhibitor (MedChemExpress, # HY-K0030) added to the PBS used for collection for the acetylation mark analyses. The whole experiment was repeated 3 times independently.

For the differentiation experiments and RNA sequencing, cells were treated as proliferative NSPCs as described above. After 5 days of culture with or without treatment with L-Carnitine or SAM, cells were split and plated on Poly-L-ornithine and laminin-coated 6-well plates, 300’000 cells per well, in proliferation medium containing 5 mg/ml of heparin, but without L-Carnitine or SAM. The next day, differentiation induction was performed by changing the medium to differentiation medium (same as proliferation medium, but containing reduced amount of growth factors, 4 ng/μl of EGF and FGFb) and 5 mg/ml of heparin. After two days, medium was changed to differentation medium containung no growth factors and 5 mg/ml heparin. Cells were collected at d0 (just before differentiation induction), d2, d4 and d6 after differentiation induction. For cell collection, the old medium was removed, and cells were washed with 2ml warm PBS. PBS was completely removed and the plates with the adherent cells were immediately frozen on dry ice and stored at - 80°C.

### Immunocytochemical staining

NSPCs were fixed with 4% PFA for 15-20 min. Cells were washed 2x for 10 min with PBS and stored at 4°C prior to staining. Cells were blocked for 45 min with blocking buffer containing 0.2% Triton-X (#X100, Sigma), 3% Donkey serum (#S30, Sigma) in Tris-buffered saline (TBS, 50 mM Tris-Cl, pH 7.4, 150 mM NaCl) and incubated with the indicated primary antibodies diluted in blocking buffer at 4°C overnight: mouse anti-Nestin (1:500, Fischer Scientific 14-5843-82), goat-anti SOX2 (1:500, R&D, AF2018-SP). Cells were washed 3x in TBS for 10 min and incubated with secondary antibodies in blocking buffer for at least 1h at room temperature, protected from light (AlexaFluor; donkey anti-mouse, donkey anti-goat,1:250, Fischer Scientific). Cells were washed 2x 10 min with TBS. Nuclei were stained with DAPI (1:5000, #D9542, Sigma) for 5 min and washed 2x with TBS. Coverslips were mounted with a self-made polyvinyl alcohol (PVA, Sigma P8136) mounting medium with 1, 4-diazabicyclo[2,2,2]octane (DABCO, Sigma D27802). To detect EdU, manufacturer’s protocol was followed (Click-iT Plus EdU Alexa Fluor 647 Imaging Kit, # 15224959, Invitrogen)

### Image acquisition and analyses

Images were acquired either with a confocal microscope (Zeiss, LSM780) with a 40x or 63x objective, or with an epifluorescent microscope (Nikon 90i or Leica DMi8) with a 20x or 40x objective. Images were analyzed using Fiji Software (ImageJ 2.3.0). SOX2 and DAPI signal were quantified using “Analyze particles” on thresholded pictures. Nestin was quantified manually as present or absent. All images were analyzed blindly.

### Cell cycle analysis

Adherent proliferative NSPCs were fixed in ice-cold 70% ethanol for 2 hours. Pellet was spun down for 5 minutes at 300g. Supernatant was removed carefully, and pellet was washed with PBS and centrifuged another 5 minutes. Pellet was re-suspended and incubated in 1 ml Propidium Iodide (PI) staining solution containing 0.1 % (v/v) Triton X-100 (Sigma, X100-100ML), 10 µg/ml PI (Life Technologies, P3566), 100 µg/ml RNase A (Life Technologies, EN0531) for 10 minutes at 37 °C. Cell cycle profile was analyzed with a Cytoflex S Flow cytometer (BeckmanCoulter). Data were analyzed using FlowJo software

### Metabolic profiling

#### Untargeted profiling of polar metabolome

Cells were grown as described in the cell culture section and stored at -80°C until processing. Polar metabolites were extracted using 400 ul ice-cold methanol:water solution (4:1, v/v) and tubes were washed with an additional 200ul ice-cold methanol:water solution. The lysed cells were further homogenized with ceramic beads in a Precellys/Cryolys Evolution tissue homogenizer and centrifuged at 13’000 rpm for 15 min at 4°C. Supernatants were collected and evaporated to dryness. Dried metabolite pellets were resuspended in methanol/water (4:1, v/v) according to protein content.

Metabolite extracts were analyzed by ultra-high-performance liquid chromatography coupled to a quadrupole time-of-flight mass spectrometer as previously described (*82*). Following data processing, the signal intensity drift was corrected using the LOWESS algorithm on pooled QC samples (*83*). The resulting peak area was used for comparative statistical analyses. We confirmed the identity of 134 metabolites using accurate mass and retention time (AMRT), compared to an in-house database created by the analysis of pure standards in the same analytical conditions. To avoid ambiguity, certain compound identities were further cross validated by MS/MS spectral matching.

Pathway enrichment analysis was performed using the online MetaboAnalyst software (5.0).

#### Amino acids and acylcarnitines quantification

As previously described (*84*), absolute quantification of acylcarnitines (AC) was conducted using stable isotope dilution approach by the addition of 250 µl of methanol spiked with internal standard solutions (ISTD) to the sample (50uL). Data were processed using Mass Hunter Quantitative Analysis (Agilent Technologies). Peak area integration was manually curated. The concentrations of AAs and ACs were normalized for the ratio of peak area between the analyte and the ISTD, and calculated using calibration curves. The lists of quantified metabolites are found in Suppl. Table 1.

### RNA-seq

RNA was extracted using RNeasy mini kit (Qiagen) according to the manufacturer’s protocol. RNA-seq data and gene counts matrices were generated by Alithea Genomics SA (Switzerland) using the BRB-seq technology (*58*).

Differential gene expression analysis between conditions was determined using the R package DESeq2 (*85*) after pre-filtering genes with low counts (rowSums > 10), with significance cut-offs set at FDR < 0.05 and fold change cut-offs set as indicated in figure legends. The sex of the samples was included as a covariate in the design of the pairwise analysis when comparing the DG and SVZ conditions. Output data were plotted with the ggplot2 R package.

For the metabolic pathway enrichment analysis, lists of genes associated with different metabolic pathways was obtained from the KEGG and REACTOME databases (*86*, *87*). For the cell type enrichment analysis, lists of marker genes for different brain cell types was obtained from a published scRNA-seq dataset (*55*). To determine enrichments from lists of DEGs we used Fisher’s exact test with significance cut-offs set at adjusted p-value < 0.01 using the Benjamini & Hochberg multiple test correction.

### ATAC-seq

ATAC-seq libraries were prepared as previously described (*88*, *89*). DG and SVZ neurospheres were dissociated in single cells. Sixty-thousand cells were re-suspended in 50 µl RSB buffer (10 mM Tris-HCl, pH 7.4, 10 mM NaCl, 3 mM MgCl2) containing 0.1% NP40 and 0.1% Tween20 and incubated on ice for 3 minutes. Lysates were washed with RSB buffer containing 0.1% Tween20 and spun down 750g for 10 min at 4 °C. Pellets were re-suspended in 50 µl transposition mix (#20034197, Illumina) and incubated for 30 min at 37 °C. Libraries were amplified by PCR with barcoded Nextera primers and sequenced.

Analysis was performed as previously described (*90*). In brief, sequencing reads were trimmed using cutadapt 3.5. Reads were aligned to the mm10 genome using bowtie2 v2.4.4. Alignments were filtered for mapping, PCR duplicates and quality using samtools v1.12. ATAC peaks were called using MACS2 v2.7.1. Peaks from all replicates were merged using bedtools v2.30, filtered against the mouse blacklist, and the numbers of reads from each replicate overlapping with the peak set was counted using featureCounts v2.0.3. Differential ATAC peak calls were determined using the R software DESeq2 after pre-filtering peaks with low counts (rowSums > 100), with significance cut-offs set at FDR < 0.05 and fold change > 1.5 in either direction. The sex of the samples was included as a covariate in the design of the pairwise analysis when comparing the DG and SVZ conditions. Output data were plotted with ggplot2. For IGV genome browser snapshots, average of replicates genome coverage files (bigwig) was generated with deepTools v3.5.2 (bamCoverage and bigwigAverage for merging replicates). Genomic features of ATAC peaks were determined using the R software ChIPSeeker v1.32.1. deepTools v3.5.2 was used to produce heat map of read density across ATAC-seq peaks. GREAT software was used to annotate ATAC peaks with genes to determine gene ontology terms associated with changes in accessibility. SEA from the MEME suite was used to identify TF motifs enriched in ATAC peaks.

### Western blot

Cell pellets were lysed in RIPA lysis buffer (10 mM Tris-HCl pH8, 150 mM NaCl, 1 mM EDTA pH8, 0.5 mM EGTA pH8, 1% NP-40, 0.5% DOC, 1% SDS) for 30 min at 4 °C.

Lysates were centrifuged at 4 °C, 13.000 rcf for 10 min and supernatant transferred to new tubes. Protein concentration was determined via BCA protein assay (Pierce, 23225), according to the manufacturer’s instructions. Samples were mixed with 1x Loading buffer (stock at 5x: 10% SDS, 0.2 M Tris HCl pH6.8, 30% Glycerol, traces of Bromophenol blue), 20x DTT (1 M) and boiled for 5 min at 95 °C. Samples were loaded onto 4-15% Mini-protean TBX gel (BioRad, 4561084) and run for 30 min at 95V then 1 h at 120 V in Running buffer (0.0125 M Tris, 0.096 M Glycine). Transfer was conducted using the Trans-Blot Turbo RTA kit (BioRad, 1704270) according to the manufacturer’s instructions. Membranes were stained with Ponceau-S solution for total protein stain (0.1% Ponceau red dye, 5% Acetic acid). Next, membranes were blocked for 1 h in 5% Milk in TBS-T (0.02 M Tris, 0.15 M NaCl, pH7.6 adjusted with HCl, 0.1% Tween20). Primary Antibody was added in appropriate dilutions in 5% Milk in TBS-T and incubated over night at 4 °C. Membranes were washed three times for 10 min in TBS-T, before incubation with secondary antibody, diluted 1:10’000 in 5% Milk in TBS-T for 1 h at RT. Membranes were again washed three times for 10 min in TBS-T. Membranes were imaged on an Odyssey DLx infrared imager (LICORbio). Western blot quantification was done using Empiria studio software (LICORbio). The following antibodies were used:

#### Primary antibodies

| Antibody | Company | Reference | Application | Dilution |
| --- | --- | --- | --- | --- |
| H3 | Abcam | ab24834 | Western blot | 1:5'000 |
| H3ac | Active Motif | 39140 | Western blot | 1:2'000 |
| H3K27me3 | Active Motif | 39055 | Western blot | 1:2'000 |
| H3K9me3 | Abcam | ab8898 | Western blot | 1:2'000 |
| H3K27ac | Abcam | ab4729 | Western blot | 1:2'000 |
| H3K9ac | Active Motif | 39138 | Western blot | 1:2'000 |
| H3K4me3 | Millipore | 05745r | Western blot | 1:2'000 |

#### Secondary antibodies

| **Antibody** | **Company** | **Reference** | **Application** | **Dilution** |
| --- | --- | --- | --- | --- |
| mouse IgG 680 | Licor | 925-68070 | Western blot | 1:10.000 |
| rabbit IgG 800 | Licor | 925-32211 | Western blot | 1:10.000 |
| mouse IgG 800 | Licor | 925-32210 | Western blot | 1:10.000 |
| rabbit IgG 680 | Licor | 925-68071 | Western blot | 1:10.000 |

### Statistical analyses

Statistical analyses were performed either with Prism (GraphPad) or R programming language (Rstudio) as following: For comparing the mean of two groups, an unpaired student t-test was used. For large data comparisons (RNA seq and Metabolomics), p-values were corrected for multiple comparisons and adjusted p-values were used. For comparing treatment effects in Western blots, one-sample t-tests were used, as each blot was normalized to its own control, set to 100%, thus all the control values were 100%. For comparing treatment effects in RNA-seq data on individual genes, 2-Way ANOVA was used, with treatment and cell type as variables, followed by posthoc tests comparing cell type differences within the same treatment group. Holm-Sidak was used for multiple comparison corrections.

The nature of the sampling (“n”), the statistical test and the p-values are described in each Figure legend. A minimum of n=3 is used for each statistical comparison. Significance was considered for p-value < 0.05.

### Data availability

The next-generation sequencing data generated in this study will be deposited on the GEO server. The metabolomics data will be deposited in the MetaboLights database.

---

**Supplementary Figure 1:**
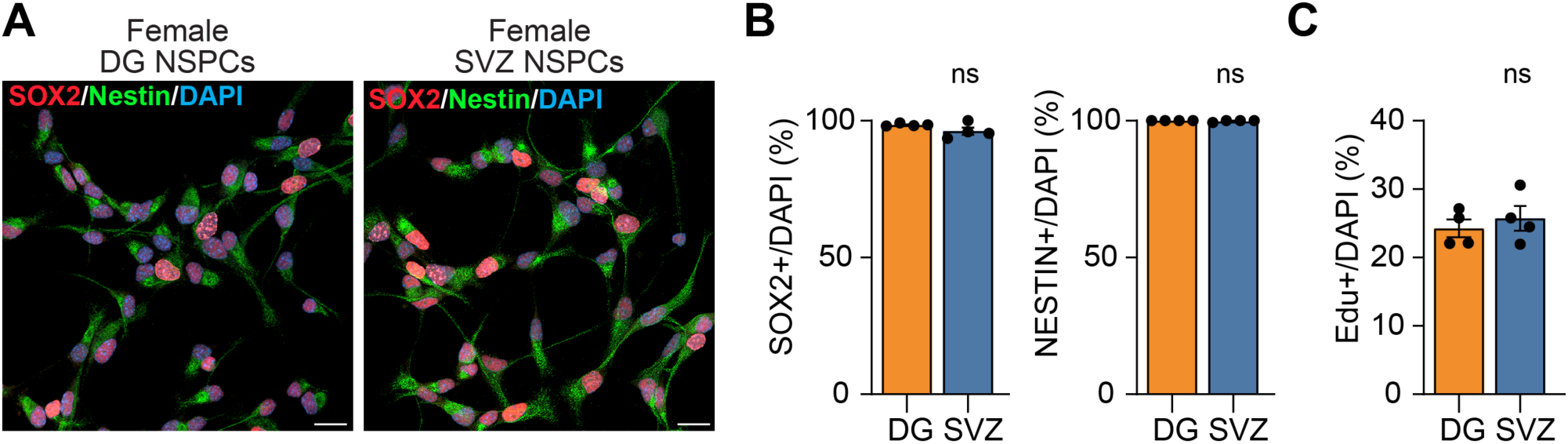
Female DG and SVZ NSPCs harbor a similar proliferation profile. **A)** Immunofluorescence staining for SOX2 and NESTIN in female DG and SVZ NSPCs. Scale bar: 20 µm. **B)** Quantification of SOX2+ and NESTIN+ cells in female DG and SVZ NSPCs. Data show individual replicates and mean ± SEM. N= 4 biological replicates, unpaired Student t-test, ns. **C)** Quantification of EdU+ cells in female DG and SVZ NSPCs. Data show individual replicates and mean ± SEM. N= 4 biological replicates, unpaired t-test, ns.

**Supplementary Figure 2:**
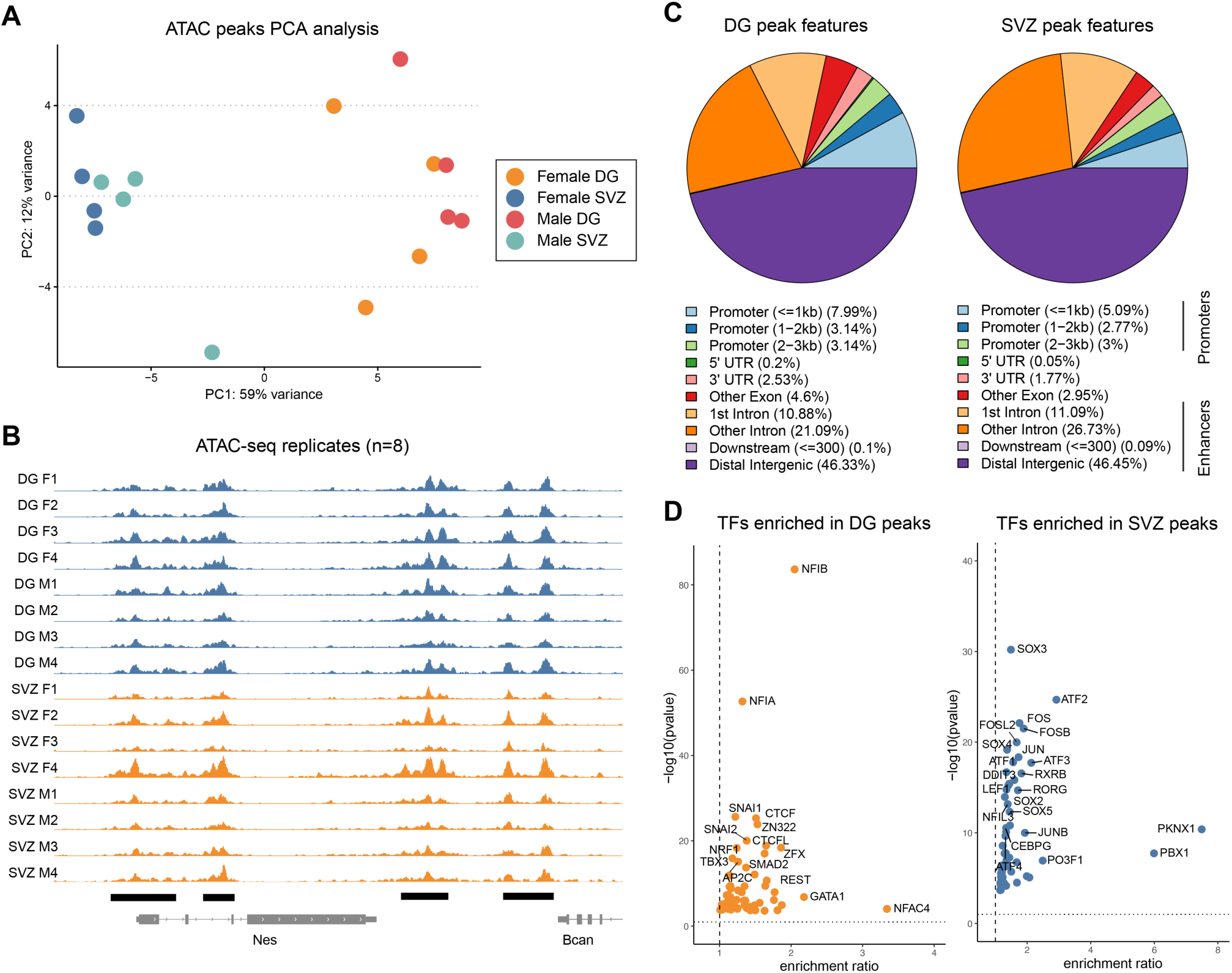
ATAC-seq analysis of DG and SVZ NSPCs predicts the type of neurons they generate. **A)** PCA of ATAC-seq from DG and SVZ NSPCs shown in the main Figure 2B, here displaying the male or female origin of samples with different colors. N= 8 biological replicates. **B)** Genome browser tracks of ATAC-seq peaks at the *Nestin* locus for all replicates of DG NSPCs (orange) and SVZ NSPCs (blue). N = 8 biological replicates, individual sample bigwig file displayed. **C)** Pie charts plot shows ATAC-seq peaks detected in DG and SVZ NSPCs within indicated genomic features. In both cell-types, ATAC-seq peaks are enriched at enhancer elements (intergenic and intronic shown in purple and orange shades). **D)** Transcription factor (TF) motif analysis of DG-specific and SVZ-specific ATAC peaks in NSPCs. Different TF repertoires bind accessible sites in DG NSPCs (NFI, SNAI, CTCF, SMAD) and SVZ NSPCs (SOX, FOS/JUN, ATF, LEF1).

**Supplementary Figure 3:**
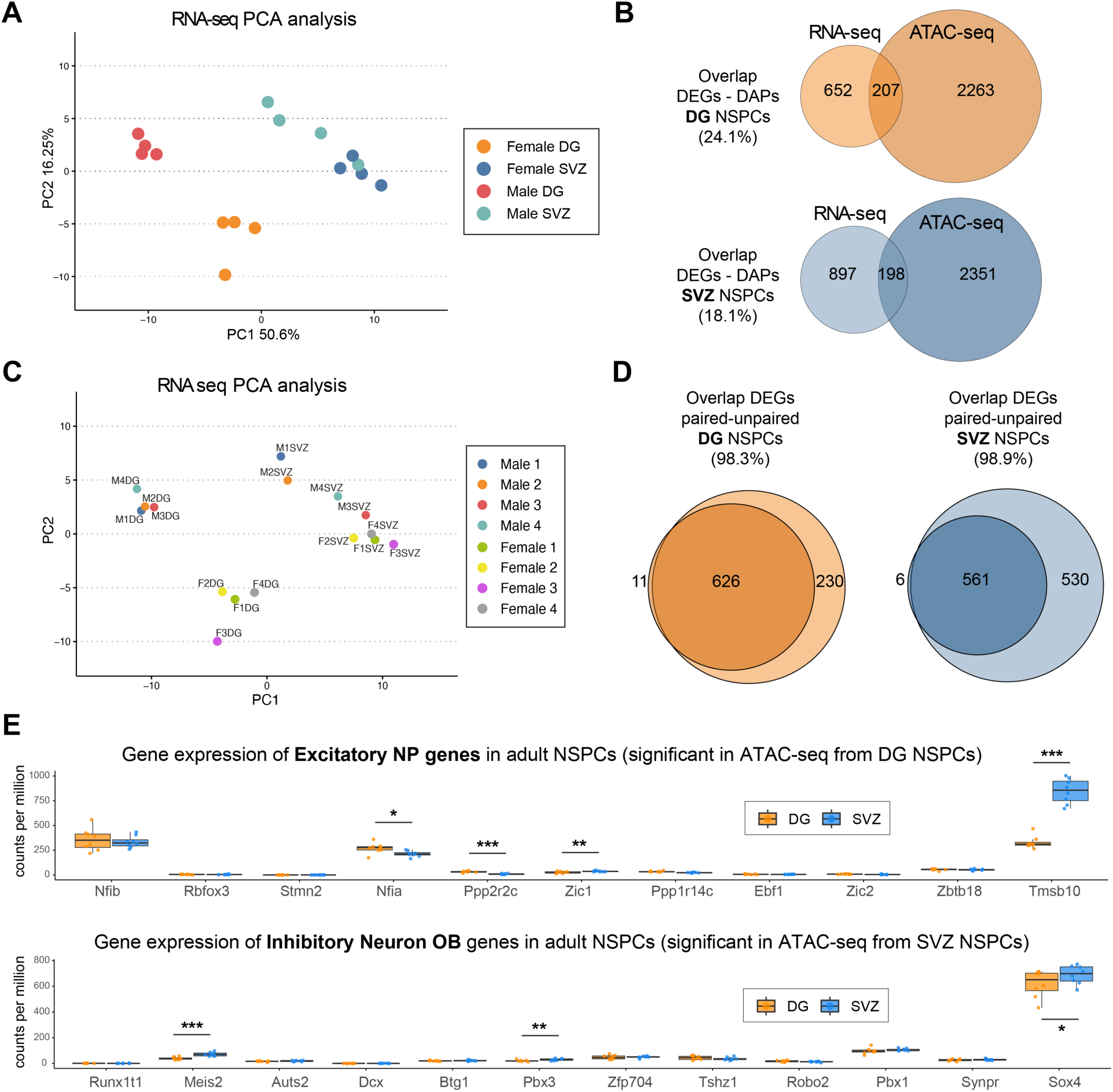
RNA-seq analysis of DG and SVZ NSPCs reveals no changes in expression of neuronal fate priming genes. **A)** PCA of RNA-seq analysis from DG and SVZ NSPCs shown in the main Figure 3B, here displaying the male or female origin of samples. N= 8 biological replicates. **B)** Top: Venn diagram displaying the overlap between DG NSPC-specific genes (RNA-seq) and genes associated with DG NSPC-specific accessible peaks (ATAC-seq). Bottom: Venn diagram displaying the overlap between SVZ NSPC-specific genes (RNA-seq) and genes associated with SVZ NSPC-specific accessible peaks (ATAC-seq). **C)** PCA of RNA-seq from DG and SVZ NSPCs displaying the paired DG and SVZ samples from the same mouse with color codes. N= 8 biological replicates. **D)** Venn diagrams displaying the overlap between DG NSPC-specific and SVZ NSPC-specific genes when comparing paired and un-paired statistical analysis of differentially expressed genes from the RNA-seq dataset. **E)** Normalized RNA-seq counts for genes that characterize the “excitatory neuronal progenitor” and “inhibitory neuron of the olfactory bulb” cell-types, from DG NSPCs (top) and SVZ NSPCs (bottom). Note the very low expression levels of most of these genes in RNA-seq, whereas they were significantly upregulated in ATAC-seq peaks (main Figure 2F). Unpaired t-test, * p<0.05, ** p<0.01, *** p<0.001.

**Supplementary Figure 4:**
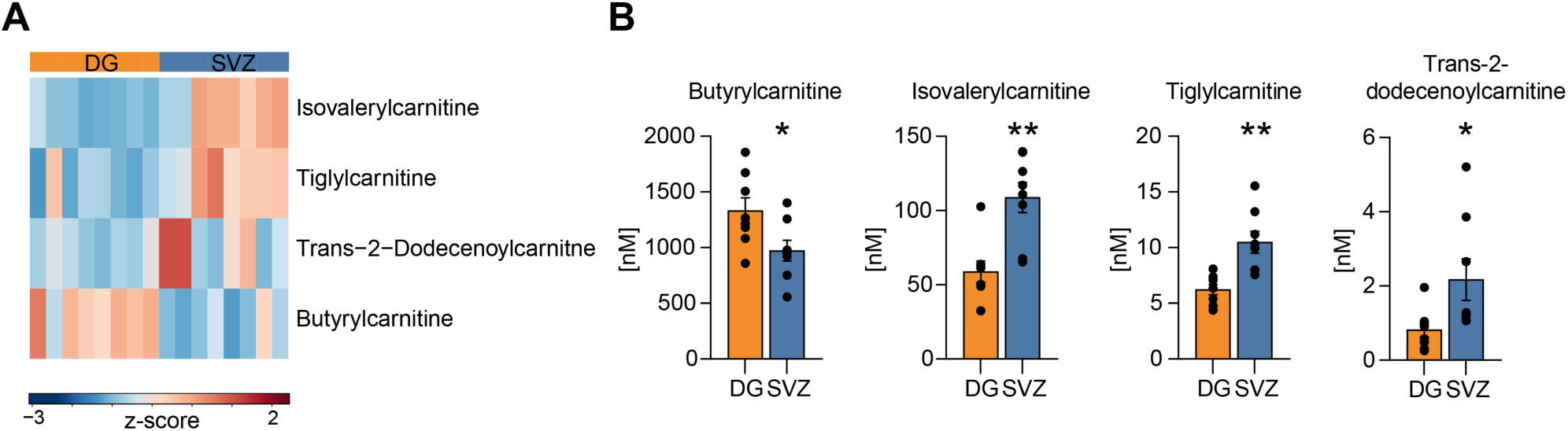
Additional acylcarnitine species which are differently abundant between DG and SVZ NSPCs. **A)** Heatmap of the additional significantly changed acylcarnitines in DG and SVZ NSPCs, p-value=0.05. **B)** Quantification of significantly changed acylcarnitines. Data show individual replicates and mean ± SEM. N= 8 biological replicates. Unpaired t-test, * p<0.05, ** p<0.01, *** p<0.001.

**Supplementary Figure 5:**
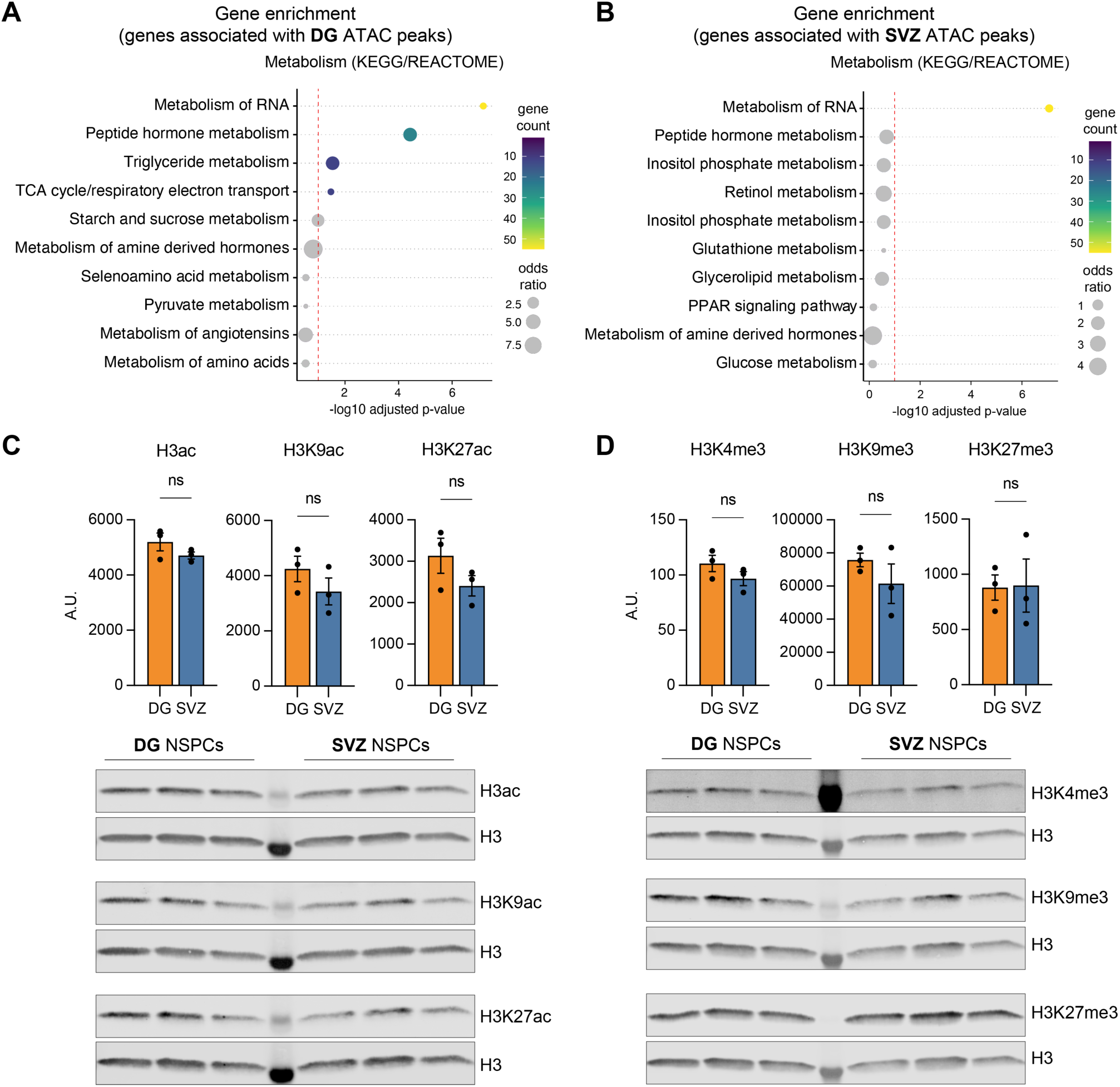
Supplementation with carnitine or SAM alters histone acetylation and methylation levels in DG and SVZ NSPCs. **A)** KEGG/REACTOM pathway enrichment analysis of the upregulated ATAC peaks in DG NSPCs, red dotted line represents p=0.05. **B)** KEGG/REACTOM pathway enrichment analysis of the upregulated ATAC peaks in SVZ NSPCs, red dotted line represents p=0.05. **C)** Western blot analysis of histone acetylation (H3ac/H3K27ac/H3K9ac) levels normalized to total histone H3 levels in DG and SVZ NSPCs. N = 3 samples per condition. Unpaired t-test, ns, non-significant. **D)** Western blot analysis of histone methylation (H3K27me3/H3K9me3/H3K4me3) levels normalized to total histone H3 levels in DG and SVZ NSPCs. N = 3 samples per condition. Unpaired t-test, ns, non-significant.

**Supplementary Figure 6:**
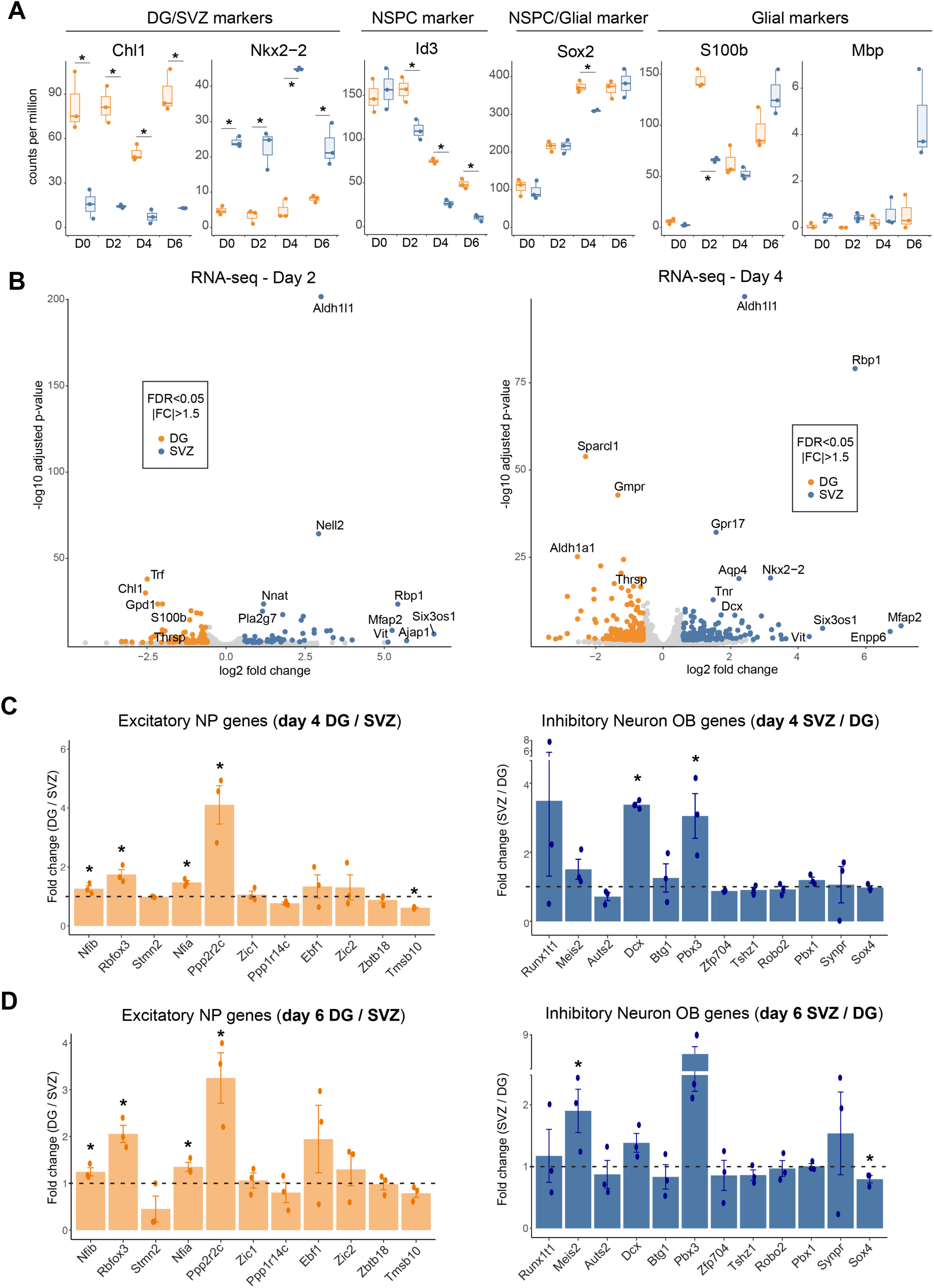
The regional identity of DG and SVZ NSPCs is maintained during differentiation. **A)** Normalized RNA-seq counts for DG and SVZ NSPCs across indicated differentiation timepoints for marker genes differentially expressed in DG/SVZ NSPCs (*Chl1, Six3*), general NSPC marker genes (*Id3, Sox2)* and glial marker genes (*S100b, Mbp*). N= 3 biological replicates. **B)** Volcano plots of differential gene expression between DG and SVZ NSPCs at Day 2 and Day 4 of differentiation. FDR < 0.05 and FC > |1.5|. N = 3 biological replicates. **C)** Graphs represent fold-change in gene expression between DG and SVZ cells at day 4 for genes that characterize the “excitatory neuronal progenitor” cell-type (left) and the “inhibitory neuron of the olfactory bulb” cell-type (right). N = 3 biological replicates. Unpaired t-test, * p<0.05. **D)** Graphs represent fold-change in gene expression between DG and SVZ cells at day 6 for genes that characterize the “excitatory neuronal progenitor” cell-type (left) and the “inhibitory neuron of the olfactory bulb” cell-type (right). N = 3 biological replicates. Unpaired t-test, * p<0.05.

**Supplementary Figure 7:**
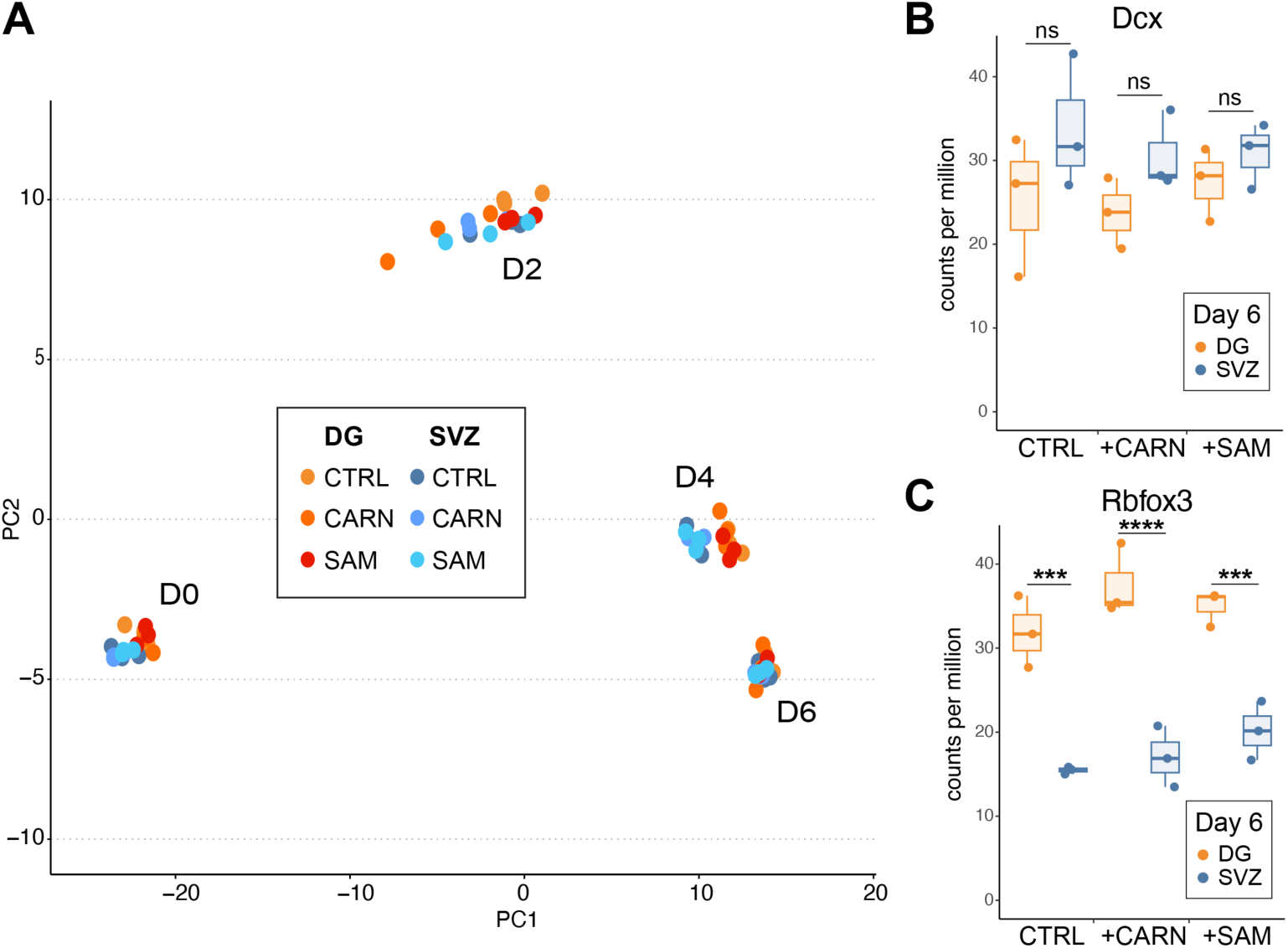
Carnitine and SAM treatment of NSPCs influences expression of neuronal-identity genes during differentiation. **A)** PCA of RNA-seq analysis from DG and SVZ NSPCs in control (CTRL), carnitine (CARN) and S-adenosylmethionine (SAM) treated conditions at all timepoints during differentiation. N = 3 biological replicates. **B)** Normalized RNA-seq counts for DG and SVZ NSPCs at day 6 of differentiation in CTRL, CARN and SAM treated conditions for the immature neuronal marker doublecortin (*Dcx*). N = 3 biological replicates per timepoint, cell type and treatment. Unpaired t-test, ns, non-significant. **C)** Normalized RNA-seq counts for DG and SVZ NSPCs at day 6 of differentiation in CTRL, CARN and SAM treated conditions for the neuronal marker *Rbfox3*. N = 3 biological replicates per timepoint, cell type and treatment. Unpaired t-test, * p<0.05, ** p<0.01, *** p<0.001.

## Notes

### Competing Interest Statement

The authors have declared no competing interest.

